# From short to long reads: enhanced protist diversity profiling via Nanopore metabarcoding

**DOI:** 10.1101/2025.06.17.660212

**Authors:** Małgorzata Chwalińska, Michał Karlicki, Sarah Romac, Fabrice Not, Anna Karnkowska

## Abstract

In the last decades environmental metabarcoding has revolutionised biodiversity research, particularly for microbial organisms such as protists, enabling large-scale assessments of diversity and ecological patterns across time and space. With the advent of long-read sequencing, Nanopore-based metabarcoding represents a promising alternative to short-read approaches. Due to the limited number of available studies, the effectiveness of Nanopore sequencing - alone or in combination with short-read data - for assessing the biodiversity and ecological patterns of protists in different ecosystems is not yet sufficiently explored. Here we present BaNaNA (Barcoding Nanopore Neat Annotator), a pipeline designed to generate high-quality OTUs and abundance estimates from Nanopore sequencing data. The performance of the pipeline was evaluated using a mock community as well as on marine and freshwater environmental samples to demonstrate its relevance for protist biodiversity and ecological studies. Our results show that BaNaNA generates high-quality full-length 18S rDNA OTUs from Nanopore long reads that are directly comparable to short-read V4-18S rDNA ASVs, supporting their synergistic use in long-term biodiversity studies. While both approaches reveal similar overall community diversity, long-read OTUs provide greater taxonomic resolution, richer phylogenetic information enabling the discovery of new clades, and yield fewer false positives. These advantages make long-read Nanopore metabarcoding not only a powerful complement but also a reliable replacement to short-read methods. By providing a pipeline for processing Nanopore data, BaNaNA paves the way for a broader application of long-read Nanopore sequencing in protist ecology and biodiversity research.

## Introduction

Microbial eukaryotes (*i*.*e*. protists) are very diverse and represent a significant part of microbial communities in the environments. Yet, their small size and limited culturing success have hampered efforts to fully explore their diversity so far. The advent of High-Throughput Sequencing (HTS) technologies applied to environmental DNA (eDNA) has revolutionized protist diversity studies for both ecological and evolutionary research (Burki et al., 2021). Metabarcoding, the most widely used approach to study protist diversity in the environment, typically targets the variable regions V4 and V9 of the 18S ribosomal RNA gene (rDNA), providing insights into the taxonomic composition and dynamics of protists over time and space (De Vargas et al., 2015; Karlicki et al., 2024; Mahé et al., 2017). The V9 region was initially preferred for protist diversity studies because it is shorter and therefore easier to sequence (Amaral-Zettler et al., 2009). However, with the development of the Illumina sequencing technology, the V4 region which offers a longer sequence with a greater variability and thus taxonomic resolution, has become the region of choice and is currently better represented in reference databases (Vaulot et al., 2022). The recent advent of long-read sequencing technologies such as PacBio or Oxford Nanopore Technology (ONT) provides longer amplicon (*i*.*e*. metabarcodes) and are being proposed as a more effective approach for microbial diversity studies. For prokaryotes, the use of full-length 16S rDNA reference sequences and even whole rDNA operons is becoming standard practice (Callahan et al., 2019; Lemoinne et al., 2024; Olivier et al., 2023; Szoboszlay et al., 2023). However, only a handful of studies have applied such long read approach to sequence the rDNA operon of protists from environmental samples (Bludau et al., 2025; Jamy et al., 2020; Overgaard et al., 2024).

Long amplicons share certain limitations with short ones, such as primer bias (Vaulot et al., 2022). Moreover, as fragment length increases, amplification efficiency often declines, making the selection of primers for long-read metabarcoding difficult (Latz et al., 2022; Sandin et al., 2022). Sequencing technology specific limitations further complicate long-read applications: PacBio sequencing, while highly accurate, remains costly and is typically limited to specialized facilities, while Nanopore is more affordable and widely accessible but has a higher error rate. More specifically, Nanopore sequencing results in a high number of indels, which impacts the generation of high-quality Molecular Operational Taxonomic Units (MOTUs). Depending on the bioinformatic processing approach applied to high-throughput sequencing (HTS) data, two main types of MOTUs can be produced: Operational Taxonomic Units (OTUs) (Edgar, 2017) and Amplicon Sequence Variants (ASVs) (Callahan et al., 2017). For short-read sequencing, denoising approaches are commonly applied to generate ASVs, which provide a high taxonomic resolution of the MOTUs present in a sample. Similar denoising strategies have also been adapted for PacBio amplicons, where low error rates allow for reliable error correction and ASV inference (Callahan et al., 2019). However, this approach is not adapted for Nanopore data due to its higher error rates and less predictable error profiles which hinder accurate error modelling. As a result, clustering reads into OTUs based on sequence similarity appears as a more appropriate strategy for MOTUs generation from Nanopore long reads (Santos et al., 2020). To ensure the reliability and reproducibility of Nanopore-based metabarcoding, there is a clear need for a standardized pipeline to process Nanopore long reads. Efforts in this direction have already been made, particularly for bacterial communities. Pipelines such as the EPI2ME platform (Oxford Nanopore Technologies), Emu (Curry et al., 2022), and MeTaPONT (Ammer-Herrmenau et al., 2021) have been developed to generate high-quality amplicons from 16S rDNA long reads. However, these approaches rely on high quality reference-based alignments which demand more comprehensive databases than those currently available for protists. Most studies applying Nanopore metabarcoding to protists have either lacked specific pipelines for clustering reads into OTUs (Gaonkar & Campbell, 2024; Hooper et al., 2023; Sandin et al., 2022) or have focused on pipelines developed for samples with low taxonomic complexity, such as clinical samples (Huggins et al., 2024; Ohta et al., 2023). NanoClust (Rodríguez-Pérez et al., 2021) has been shown to be effective for low-complexity protist communities (Huggins et al., 2024), but so far only Natrix2 (Deep et al., 2023) has been shown to properly process long-read metabarcoding data for protists (Bludau et al., 2025). In addition to developing robust methods for generating OTUs from Nanopore long-read amplicons (Overgaard et al., 2024), a major challenge is to accurately estimate OTUs abundances.

Longer meta-barcodes offer significant advantages as they provide higher taxonomic resolution and therefore allow a more detailed understanding of protist communities, from detailed biogeography to evolutionary studies (Gaonkar & Campbell, 2024; Jamy et al., 2020, 2022). However, the extensive datasets generated up to now from short amplicons remain an invaluable resource, especially for large-scale (*e*.*g*. De Vargas et al., 2015) or long-term studies (*e*.*g*. Yeh & Fuhrman, 2022). It is therefore important to evaluate how these approaches complement each other and can eventually be combined. While some attempts have been made for groups of organisms such as zooplankton (Chang et al., 2024), little is known about the impacts on ecological analyses or the feasibility of integrating short and long amplicons in comparative studies for protists. A recent comparison of Illumina V9-18S rDNA and Nanopore 18S rDNA protist metabarcodes from sediment samples (Bludau et al., 2025) demonstrated that Nanopore metabarcodes provide higher taxonomic resolution for protists, while both methods revealed similar diversity and basic community patterns.

Here, we further addressed these critical challenges by analysing both Illumina and Nanopore metabarcoding data from a protist mock community, as well as from distinct environmental sample sets representing marine and freshwater ecosystems. To generate high-quality OTUs from Nanopore data, we introduce the BaNaNA pipeline, primarily designed for metabarcoding analysis of microbial eukaryotes but suitable for other taxa as well. We evaluated the effectiveness of Nanopore long-read 18S rDNA and Illumina short-read V4-18S rDNA metabarcoding, providing a detailed assessment of how each method influences biodiversity and ecological interpretations, ultimately emphasizing the advantages of Nanopore-based metabarcoding and the potential for combining both approaches.

## Methods

### Samples collection

Freshwater samples (Table S1) were collected at the end of July to the beginning of August in year 2020 from five lakes in the Great Masurian Lakeland District in north-eastern Poland. Samples were collected with a modified Bernatowicz sampler from photic and aphotic zones of each lake, prefiltered through 150 µm mesh-size net to remove zooplankton and bigger particles and then filtered under pressure through 0.2 µm membrane Nucleopore filters (Whatman, Maidstone, UK). Filters were frozen at -20°C and kept in a -80°C freezer for long-time storage.

Marine samples (Table S2) were collected during one of the annual MOOSE-GE campaigns (https://campagnes.flotteoceanographique.fr/campagnes/17001500/) (Coppola et al., 2019) from 31 August to 23 September 2017. A volume of 20 L of seawater from 2 x 12L-bottles was taken at the surface, DCM and deep waters (ca 2000m depth). The water was prefiltered through 180 µm and filtered through 0.2 and 3 μm, 47 mm Nucleopore polycarbonate filters (Whatman, Maidstone, UK). After filtration, filters from both size fractions were flash-frozen in liquid nitrogen and stored independently at -80°C until DNA extraction.

### Mock community preparation

The mock community was composed of seven species from culture collections (Table S3). Species represent six groups of protists: Haptophyta, Euglenozoa, Chlorophyta, Ciliophora, Dinoflagellata and Cryptophyta (2 species). Among these, *Prymnesium parvum* (Haptophyta), *Euglena gracilis* (Euglenozoa) and *Chlorella variabilis* (Chlorophyta) were highly abundant, while *Paramecium bursaria* (Ciliophora), *Gymnodinium fuscum* (Dinoflagellata), *Cryptomonas paramecium* (Cryptophyta) and *Cryptomonas gyropyrenoidosa* (Cryptophyta) were added at low concentration (Table S3). In addition to the taxonomic differences, the species within the mock community differed considerably in cell size, morphology and cell number designed to mimic natural samples.

Cells abundances were manually calculated using a Fuchs-Rosenthal chamber, all species were combined and filtered under pressure through 0.2 μm membrane Nucleopore filters and frozen at -80°C. For *Cryptomonas gyropyrenoidosa* which did not have 18S rDNA reference sequence available, we isolated DNA from the culture using NucleoSpin Tissue XS kit and amplified 18S rDNA gene using SA (5’ AACCTGGTTGATCCTGCCAGT 3’) (Medlin et al., 1988) and EukB (5’ TGATCCTTCTGCAGGTTCACCTAC 3’) (Medlin et al., 1988) primers (Supplementary Materials), purified the DNA using PCR Mini Kit (Syngen) and sequenced using Sanger with additional primer Euk528F (5’ CGGTAATTCCAGCTCC 3’) (Edgcomb et al., 2011). Fragments were then assembled using Lasergene Seqman Pro.

### DNA isolation and amplification

DNA from freshwater samples and mock community was extracted from ¼of the filter using the GeneMATRIX Soil DNA Purification Kit (EURx), its concentration was measured using NanoDrop (Thermo Scientific) and frozen at - 80°C. DNA from marine samples were extracted using a modified protocol from the NucleoSpin Plant II Mini or Midi kits (Macherey-Nagel), depending on the planktonic size-fraction. Detailed DNA extraction protocol is available on the online protocol repository protocols.io : dx.doi.org/10.17504/protocols.io.kxygxy5xdl8j/v1.

DNA extracts were diluted to 5 ng/µL and amplified in three replicates for both sequencing methodologies using Phusion High-Fidelity DNA polymerase (Finnzymes; ThermoFisher). For Illumina strategy, the V4 region of the 18S rRNA gene (∼380 bp) was targeted using the primers TAReuk454FWD1 (5’ CCAGCASCYGCGGTAATTCC 3’) and TAReukREV3 (5’ ACTTTCGTTCTTGATYRA 3’) (Stoeck et al., 2010). Amplification for marine samples is detailed on protocols.io : dx.doi.org/10.17504/protocols.io.bzucp6sw, while the freshwater and mock community amplification protocol can be found in the Supplementary Materials of this article. For Nanopore sequencing, a fragment from the beginning of 18S all the way to the D2 region of 28S rDNA (amplicon size 3200 bp) was targeted using the SA (5’ TTTCTGTTGGTGCTGATATTGCAACCTGGTTGATCCTGCCAGT 3’) (Medlin et al., 1988) and D2C-R (5’ ACTTGCCTGTCGCTCTATCTTCCCTTGGTCCGTGTTTCAAGA 3’) (Scholin et al., 1994) primers extended by Nanopore adapters. The detailed protocol and the sequence of Nanopore adapters are placed in the Supplementary Materials. After PCRs, the replicates were merged and purified together using a PCR Mini Kit (Syngen).

### Illumina library preparation and sequencing

Library for freshwater samples and mock community was prepared and sequenced in the Genomics Core Facility at the Centre of New Technologies (University of Warsaw, Poland) using the Illumina MiSeq platform with 2x 250 bp. Regarding marine samples, library adapter ligation and sequencing were performed in the same conditions by Fasteris (www.fasteris.com, Plan-les-Ouates, Switzerland) on a 2 × 250 bp MiSeq Illumina.

### Nanopore library preparation and sequencing

Nanopore libraries were prepared using PCR Barcoding Expansion 1-12 (EXP-PBC001) and Ligation Sequencing Kit (SQK-LSK114). Samples were sequenced on MinION Mk1B device using R10.4.1 flow cells.

### Illumina data analysis

Quality of raw sequences was checked using FastQC v0.11.5 (Andrews, 2010). Primers were removed using Cutadapt plugin for QIIME2 2023.9.1 environment (https://github.com/qiime2/q2-cutadapt) (Bolyen et al., 2019). Representative sequences (ASVs – Amplicon Sequence Variants) were created using DADA2 (Callahan et al., 2016) in QIIME2 environment using DADA2 denoise-paired function (https://github.com/qiime2/q2-dada2). Final ASVs had their taxonomy assigned using global alignment method of VSEARCH v2.7.1 (Rognes et al., 2016) to PR2 database v5.0.0 (Guillou et al., 2012) with minimum threshold of 70% identity and minimum query coverage of 90%.

### Nanopore data analysis

To overcome the high error rate associated with Nanopore sequencing technology, we have developed the BaNaNA (Barcoding Nanopore Neat Annotator) (https://github.com/ibe-uw/BaNaNA) - a Snakemake (Mölder et al., 2021) pipeline to generate high-quality representative sequences, also known as Operational Taxonomic Units (OTUs), from long-reads amplicons (Fig. 1).

**Fig. 1.**
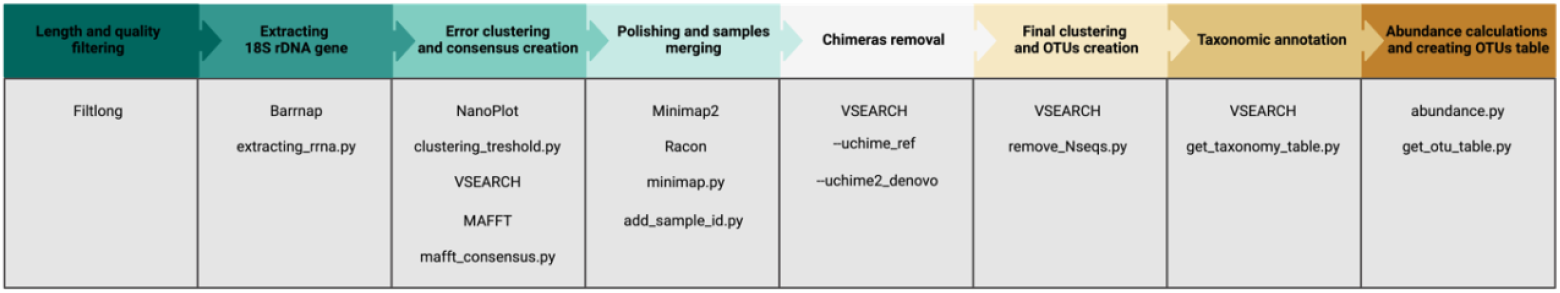
Overview of the BaNaNA pipeline for obtaining OTUs from Nanopore reads. The diagram illustrates the sequential steps of the BaNaNA workflow, along with intergrated tools and custom scripts used at each stage of the analysis. (Created with Biorender.com)

The pipeline includes multiple steps which are briefly outlined here. First, the raw reads were basecalled using the duplex option and super-accurate model and demultiplexed with Dorado v0.5.1+a7fb3e3 (Oxford Nanopore Technologies https://github.com/nanoporetech/dorado). We then filtered the reads for length and quality using Filtlong v0.2.1 (https://github.com/rrwick/Filtlong) and extracted rDNA fragments using Barrnap 0.9 (https://github.com/tseemann/barrnap). An additional quality check of the rDNA fragments was performed using the custom Python 3.9.18 script and Biopython tool (Cock et al., 2009) (extracting_rrna.py) which checked the presence of the correct rDNA structure (18S, 5.8S, 28S) and the length of each region. For further analysis we kept only 18S rDNA gene. We then calculated the average read quality with NanoPlot 1.42.0 (De Coster & Rademakers, 2023) and FASTQ reads according to Filtlong and used this value as the threshold for the VSEARCH v2.7.1 (Rognes et al., 2016) clustering step. To obtain consensus sequences, we used the custom script mafft_consensus.py, which uses MAFFT v7.310 (Katoh & Standley, 2013) to create an alignment within clusters and compares each position to create final sequences. In the next step, we used Minimap2 2.24-r1122 (Li, 2018) and Racon 1.5.0 (Vaser et al., 2017) to polish the sequences. Next, we added the names of the samples to the sequence header to make them easier to identify. After merging all processed samples, we used VSEARCH to remove chimeric sequences. We first used the PR2 5.0.0 (Guillou et al., 2012) database as a reference and then recognised chimaeras *de novo*. Final clustering at 99% identity with VSEARCH allowed us to remove duplicate sequences and create final Operational Taxonomic Units (OTUs). Due to different lengths of sequences within clusters during consensus building and polishing, some OTUs accumulated a large number of ambiguous bases represented as Ns, which reduces their quality; the remove_Nseqs.py script was used to remove them. Subsequently, the obtained OTUs were taxonomically annotated against the PR2 database using the global alignment method implemented in VSEARCH, using a minimum identity of 70% and a minimum query coverage of 90% as parameters. The abundances were calculated by counting the number of sequences in the clusters from the first clustering. After calculating the abundances, the OTU table is created with all samples. Additionally, the BaNaNA pipeline allows the use of various databases and modifications with respect to different primers and different fragments of the rDNA.

### Extraction of V4-tags from Nanopore sequences

To assess the impact of sequence length on taxonomic annotation and to investigate the possibility of integrating Nanopore and Illumina data, we extracted V4-tags from the Nanopore OTUs. For this, we used the same primer sequences as for Illumina sequencing as well as customised R and Python scripts with the help of Biostrings package (Pagès et al., 2021) and Biopython tool (Cock et al., 2009). The extracted V4 tags were then taxonomically assigned independently from the Nanopore OTUs using VSEARCH v2.7.1 (Rognes et al., 2016).

### Species-level classification and false positives assessment

To assess the representation of our mock community species in the PR2 database 5.0.0 (Guillou et al., 2012), we performed a global alignment of the sequences of the reference strains (Table S3) with the sequences of the database using VSEARCH v2.7.1 (Rognes et al., 2016). A sequence was classified to species level if the best hit had at least 99% similarity with the reference strain sequence or if the best hit was already classified as the same species. If the best hit was only classified to genus level and had less than 99% similarity to the strain reference, the sequence was assigned to genus level. All other hits, which were classified to different genera were classified as false positives.

### Phylogenetic tree construction from Nanopore OTUs

To construct the phylogenetic tree, we extracted freshwater and marine OTUs whose percent identity with the closest PR2 5.0.0 (Guillou et al., 2012) reference was between 80% and 97% and therefore likely represent taxa not represented in the database. Sequences were aligned using MAFFT v7.310 (Katoh & Standley, 2013) and the resulting alignment was trimmed with trimAl v1.4.rev15 (Capella-Gutiérrez et al., 2009) using the - automated1 option. Phylogenetic inference was performed using IQ-TREE 2.0.6 (Minh et al., 2020) with the - m MFP option to determine the optimal substitution model. After tree construction, we manually inspected both the alignment and the tree to identify OTUs the taxonomy of which was inconsistent with the general taxonomy of the clade in which they were placed. Such OTUs were further curated by performing BLAST (Altschul et al., 1990) searches against the NCBI nt database (Wheeler et al., 2007) to refine the taxonomy (3 OTUs) or, if necessary, remove them from the dataset (26 OTUs). The sequences were then realigned, trimmed and the tree was recalculated using the same methods as described above. The TIM2+F+R10 model was selected as the best fitting model for the final tree. The resulting tree was visualised using RStudio (RStudio Team, 2020) with the ggtree (Yu et al., 2017), treeio (Wang et al., 2020), dplyr (Wickham et al., 2022) and readxl (Wickham & Bryan, 2019) packages.

### Statistical analyses

Further statistical analyses were performed in RStudio (RStudio Team, 2020) using libraries: phyloseq (McMurdie & Holmes, 2013), vegan (Oksanen et al., 2022), tidyverse (Wickham et al., 2019), reshape2 (Wickham, 2007), readxl (Wickham & Bryan, 2019) and ggplot2 (Wickham, 2016). From all datasets we removed low abundance sequences. For environmental samples those were sequences which were present less than 5 times in the whole dataset. Due to the low complexity of the mock community, we decided to remove sequences which had relative abundance below 0.01% within any one of the samples. For beta-diversity we aggregated ASVs/OTUs at the genus level and calculated Bray-Curtis dissimilarity index combined with NMDS ordination method.

## Results

### Comparison of Illumina and Nanopore metabarcoding using a mock community

We first analysed a mock community of seven species representing major protist lineages mixed at a defined concentration (Table S3). Illumina V4-18S rRNA gene sequencing yielded 622,100 raw reads which after processing generated a total of 268 ASVs with an average length of 379 bp (Table S4). Nanopore sequencing yielded 594,076 raw reads encompassing the full 18S rRNA gene ITS1, 5.8S rRNA gene, ITS2 and part of 28S rRNA gene. After *in silico* extraction of the 18S rRNA gene fragments and OTUs clustering using the BaNaNA pipeline, we identified 147 OTUs with an average sequence length of 1,843 bp (Table S5). Further extraction of the V4 region of 18S rRNA gene (V4-tags) from the OTUs yielded 145 sequences with an average length of 416.1 bp.

Illumina V4-18S rDNA ASVs revealed a taxonomic composition that differed substantially from the original community structure of the mock community (Fig. 2A). Dinoflagellates were the most abundant group, followed by haptophytes, ciliates and cryptophytes, while euglenozoans were nearly absent from the dataset. In addition, a considerable number of Illumina ASVs were classified as “other taxa”, representing organisms that were not intentionally included in the mock community. The full-length 18S rDNA OTUs from Nanopore sequencing more closely mirrored the expected community composition, with haptophytes being the most abundant, followed by euglenozoans.

**Fig. 2.**
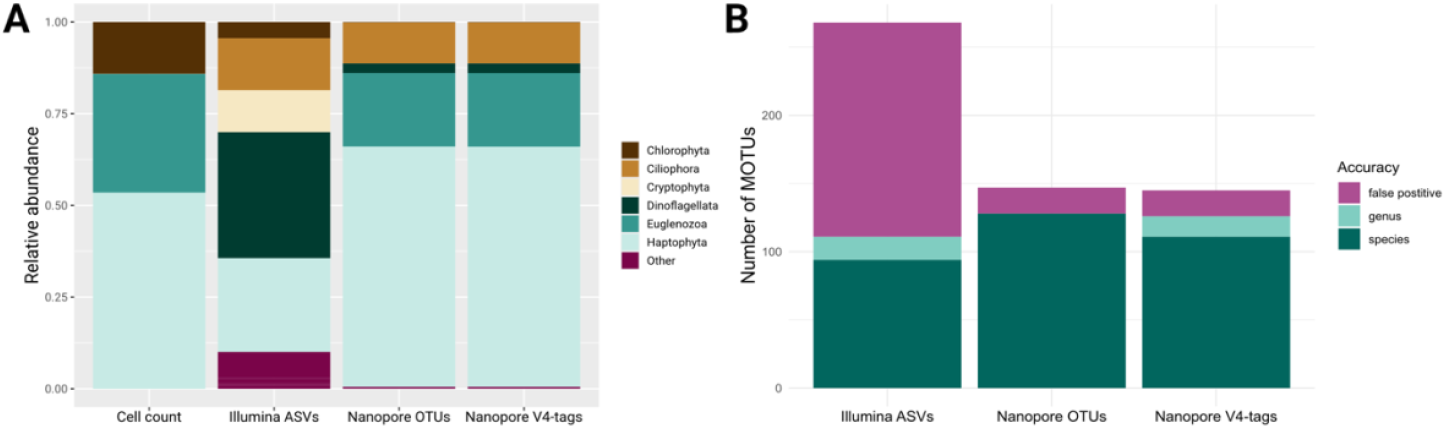
**(A)** Relative abundance of species in the mock community at the subdivision level, as determined by cell counts and all sequencing approaches. Taxa that were not intentionally included in the mock community are grouped as ‘Other’. **(B)** Accuracy of MOTUs identification in the mock community. Bar plots display the accuracy of taxonomic assignment for ASVs and OTUs derived from the mock community, based on comparisons with known species composition.

However, ciliates replaced chlorophytes as the third most abundant group. No significant differences were observed between the V4-tags and the full-length 18S rDNA OTUs. Nanopore datasets contained also additional taxa that were absent from the original mock community, but their number and diversity appeared much larger in the Illumina data (Fig. S1). A detailed analysis of the additional taxa suggests that although many of them probably represent biases of the ASVs or OTUs generation some might correspond to real occurrences of the organisms rather than sequencing errors and some other could represent minor contaminations (Fig. S2).

Ultimately, neither sequencing technology was able to accurately identify all seven taxa in the mock community at the species level or quantify them correctly (Table 1). Illumina sequencing failed to detect *Paramecium bursaria* at the species level, while Nanopore did not identify *Cryptomonas gyropyrenoidosa* and *Chlorella variabilis*. The Illumina data exhibited significant biases in taxon relative abundances, with *Euglena gracilis* and *Chlorella variabilis* being substantially underestimated by three and two orders of magnitude respectively, while *Gymnodinium fuscum* and both *Cryptomonas* species were overestimated by three and two orders of magnitude respectively. Both Nanopore (*i*.*e*. full-length 18S rDNA OTUs and V4-tags) approaches were mostly in agreement and accurately estimated the relative abundance of *Euglena gracilis* and *Cryptomonas paramecium* and overestimated *Gymnodinium fuscum* by two orders of magnitude (Table 1). However, for *Paramecium bursaria*, the two Nanopore approaches yielded different results: the full-length 18S rDNA OTUs overestimated abundance by three orders of magnitude, whereas the V4-tags provided an estimate much closer to the actual cell abundance. *Prymnesium parvum* was the only taxon for which relative abundance was consistently estimated to the same level of magnitude across sequencing technologies and matched its proportion in the mock community based on cell counts.

**Table 1.**
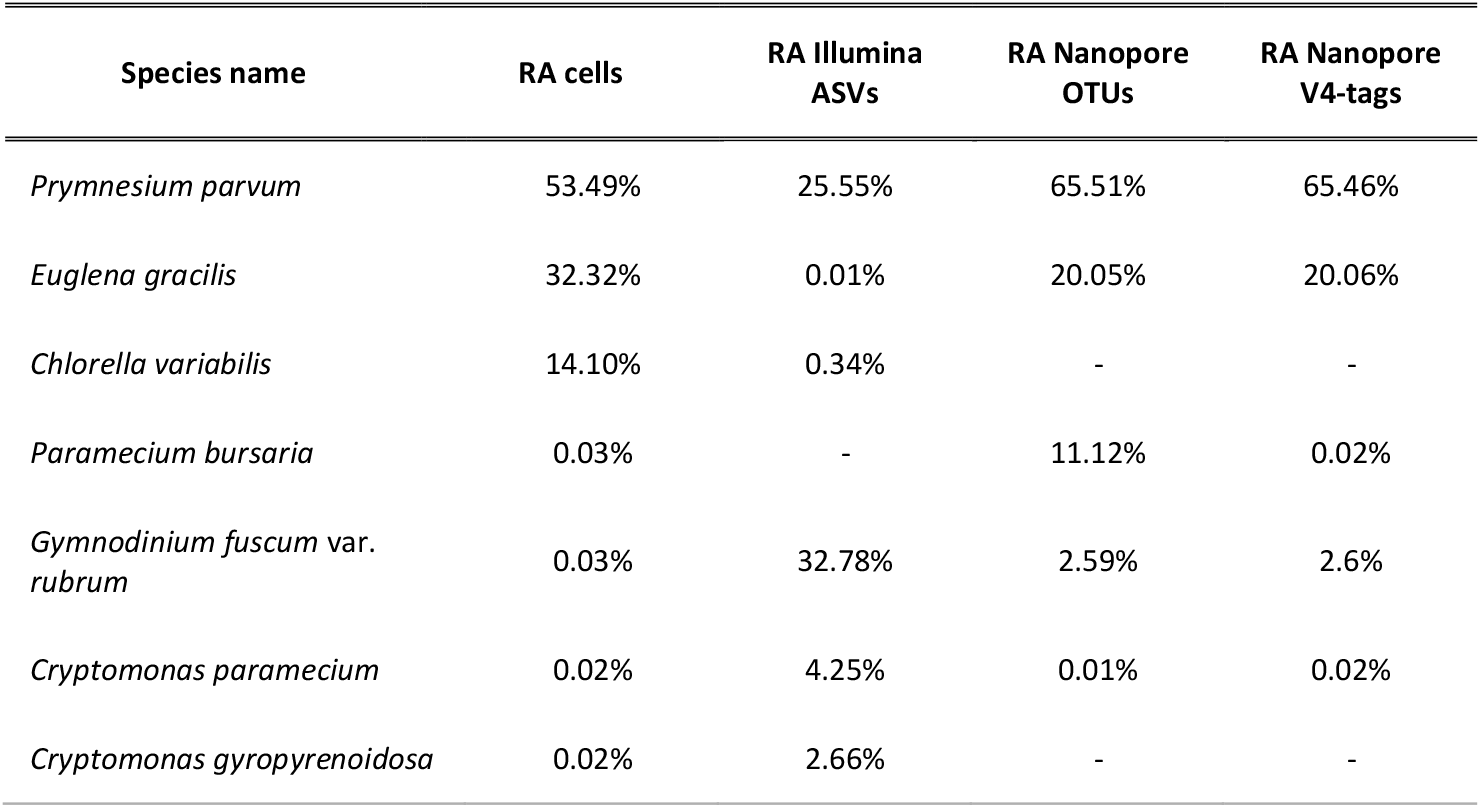
Species composition of the mock community with relative abundances (RA) based on cell count and MOTUs classified properly to the species level.

The presence of additional taxa and the poor relative abundance obtained at the species level prompted us to investigate the impact of sequencing technology on the accuracy of taxonomic annotation. The V4-18S rDNA ASVs from Illumina sequencing had the fewest representative sequences classified to species-level and the highest number of false positives (see Materials and Methods) (Fig. 2B). In contrast, full-length 18S rDNA OTUs had the highest number of species-level classifications compared to both Illumina V4-18S rDNA ASVs and V4-tags. This suggests that shorter fragments reduce the accuracy of taxonomic annotation and fail to classify some OTUs to species level, whereas longer fragments provide sufficient resolution for proper classification, as long as reference database is accurate. This is also emphasized when comparing full-length 18S rDNA OTUs with V4-tags, as both were amplified using the same primers, disregarding potential primer bias. The trend was especially pronounced in *Paramecium bursaria*, where species-level OTUs classification significantly decreased in V4-tags compared to full-length 18S rDNA OTUs (Table 1).

### Comparison of Illumina and Nanopore metabarcoding using environmental samples

To evaluate the performance of both sequencing approaches and data analysis pipelines on complex natural communities, we sequenced and analysed 10 freshwater samples from lakes and 12 marine samples. Illumina sequencing yielded from 23,132 to 95,084 raw reads per sample with average length of 250.5 bp, which after processing resulted in a total of 1,668 ASVs (average length: 376.5 bp) from lakes and 2,716 ASVs (average length: 380.6 bp) from marine samples (Table S4). Nanopore sequencing yielded from 77,813 to 591,619 raw reads per samples with an average length of 2404.3 bp. After extraction of the 18S rDNA fragments and OTUs clustering, we identified 1,068 OTUs (average length: 1,766.5 bp) from lakes and 1,230 OTUs (average length: 1,785.9 bp) from marine samples (Table S5). The extraction of V4-tags from the OTUs yielded 1,065 sequences (average length: 379.2 bp) from lakes and 1,200 sequences (average length: 384.7 bp) from marine samples.

According to rarefaction curves (Fig. 3A), all samples sequenced with Illumina reached saturation. Yet, notable differences in the numbers of species could be observed between samples, with the freshwater samples having a higher average number of species than photic marine samples (189.7 for freshwater *vs* 169.4 for surface + DCM) (Fig. S3, Fig. S4). Marine deep-sea samples exhibited the lowest average number of species (100.7) (Fig. S4). Although samples were pooled at equal concentrations during library preparation, there were significant differences in the number of raw reads obtained during Nanopore sequencing (Table S5). Nevertheless, all Nanopore samples, except the aphotic sample from Babięty Wielkie lake (BAB-APH) reached a well visible saturation plateau (Fig. S5, Fig. S6). Unlike for Illumina, in case of Nanopore OTUs marine photic samples had a slightly higher number of species than the freshwater samples (110.7 for surface + DCM vs 95.5 for freshwater). Marine deep-sea samples consistently had the lowest average number of species (average of 49.0) (Fig. S6). We observed differences in the number of recovered taxa between the two sequencing technologies at all taxonomic levels (Fig. 3B). Illumina systematically identified more taxa than Nanopore, the differences were particularly pronounced at low taxonomic levels, where Illumina contained higher average number of species per sample (166.1) than Nanopore (92.6) (Table S4, Table S5). Both technologies identified some unique taxa, not detected by the other technology.

**Fig. 3.**
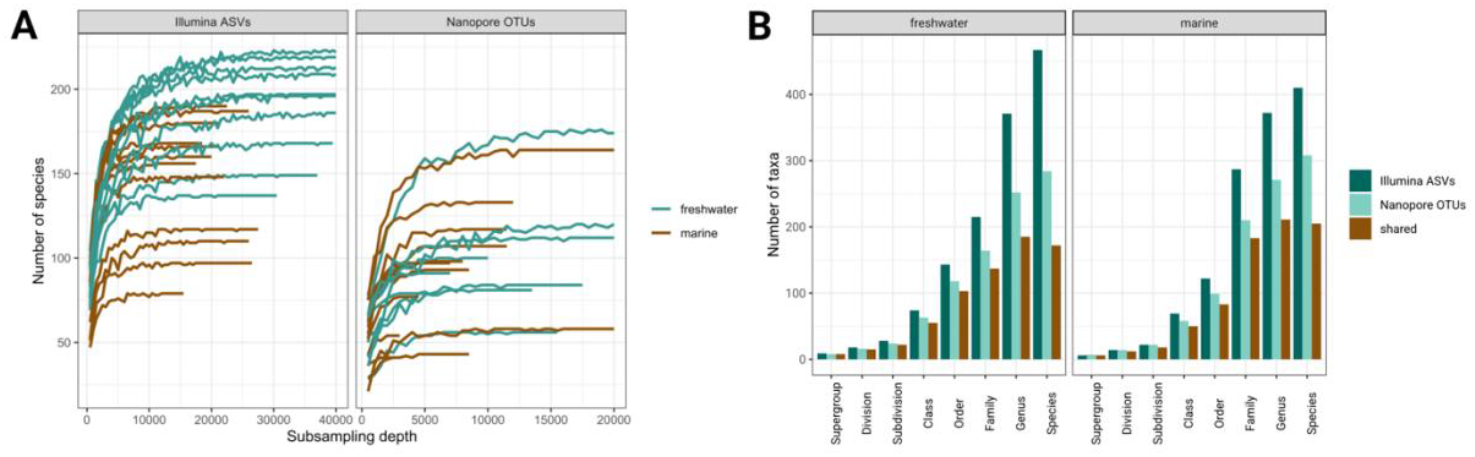
**(A)** Rarefaction curves showing the number of observed species per sample sequenced with Illumina and Nanopore technologies. **(B)** Number of taxa identified at different taxonomic ranks in freshwater and marine environmental samples using Illumina and Nanopore sequencing, along with the number of taxa shared by both datasets.

We compared the relative abundances of main protist groups between Illumina and Nanopore sequencing for both freshwater (Fig. 4A) and marine (Fig. 4B) samples. Overall, the relative abundances at higher taxonomic levels were very similar. In lakes, both methods identified dinoflagellates and perkinseans as the most abundant groups, followed by ciliates, cryptophytes and chlorophytes. Moreover, there were no major differences in the composition of the major protist groups between the photic and aphotic zones of the lakes. In the marine environment, dinoflagellates and radiolarians dominated, followed by Gyrista and Euglenozoa (Diplonemida), the latter being detected exclusively by Nanopore sequencing. Samples from the sea surface showed a lower relative abundance of radiolarians than samples from the deep chlorophyll maximum, while samples from the deep sea contained the highest proportions of radiolarians and lacked phototrophic groups such as Bigyra and Chlorophyta, which occur in the photic zone. At higher taxonomic levels, the results of the V4-tags were more similar to the full-length 18S rDNA OTUs than to the V4-18S rDNA ASVs (Fig. 4 A, B), highlighting the biases introduced by the different primer pairs for Nanopore and Illumina experiments. On the other hand, the full-length 18S rDNA OTUs generally showed lower identity to the closest reference sequences than the shorter V4-18S rDNA ASVs and V4-tags (Fig. 4C).

**Fig 4.**
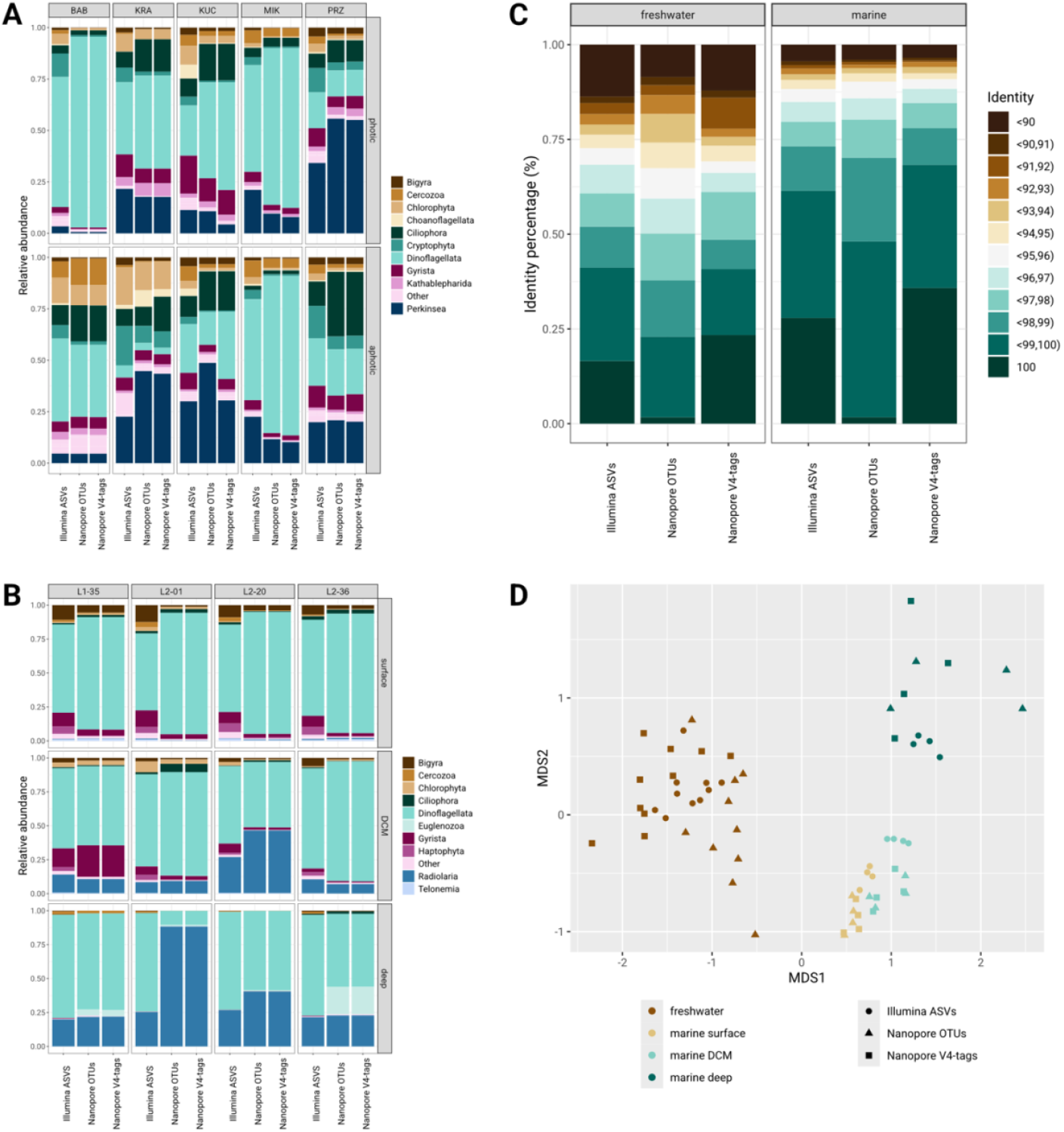
Relative abundances of the ten most prevalent subdivisions in freshwater **(A)** and marine **(B)** samples across sampling spots and depths. **(C)** Identity distribution of MOTUs sequences compared to reference sequences in the PR2 database. **(D)** NMDS ordination plot of all environmental samples based on Bray-Curtis dissimilarity of relative abundances of MOTUs grouped at the genus level.

We also observed notable differences in the proportion of sequences annotated at low identity levels (95% or less) between the different environments (Fig. 4C). In freshwater samples, the proportion of such sequences was more than twice as high as in the marine samples (32% in Illumina freshwater vs. 15% in Illumina marine; 41% in Nanopore freshwater vs. 14% in Nanopore marine). This discrepancy could be directly related to the unbalanced number of reference sequences available for freshwater and marine protists in the databases.

To further assess the impact of sequencing technologies on ecological interpretation of the data, we estimated the overall variation in genera composition between environments and calculated the Bray-Curtis beta diversity index (Fig. 4D). Despite differences in taxonomic resolution at lower levels, both sequencing approaches revealed similar microbial community structures across the analysed environments. Illumina V4-18S rDNA ASVs, full-length 18S rDNA OTUs, and V4-tags were all clustered into three distinct groups corresponding to their respective environments. This suggests that the discrepancies between sequencing technologies were primarily driven by low-abundance taxa and had minimal impact on the general ecological patterns.

### Diversity of novel OTUs obtained from Nanopore sequencing

We recovered 747 full-length 18S rDNA OTUs from freshwater and marine environments, which were considered as novel according to the sequence identities threshold from 80% to 97% to their nearest references in the PR2 reference database. The 513 novel OTUs from freshwater environments represented 48% of all obtained freshwater OTUs and 234 novel OTUs from marine environments represented 19% of all marine OTUs obtained in this study. The novel OTUs are a valuable addition to the reference databases as they represent 7.6% (freshwater) and 3.5% (marine) of all eukaryotic sequences currently present in the PR2. Availability of the full-length 18S rDNA OTUs enable the exploration of novel OTU diversity by reconstructing phylogenetic trees, allowing the identification of not only new sequences but also potentially novel clades. After manually curating the alignment, we inferred a phylogenetic tree from 721 novel OTUs (Fig. 5) which grouped into clades corresponding to six supergroups and fifteen divisions.

**Fig. 5.**
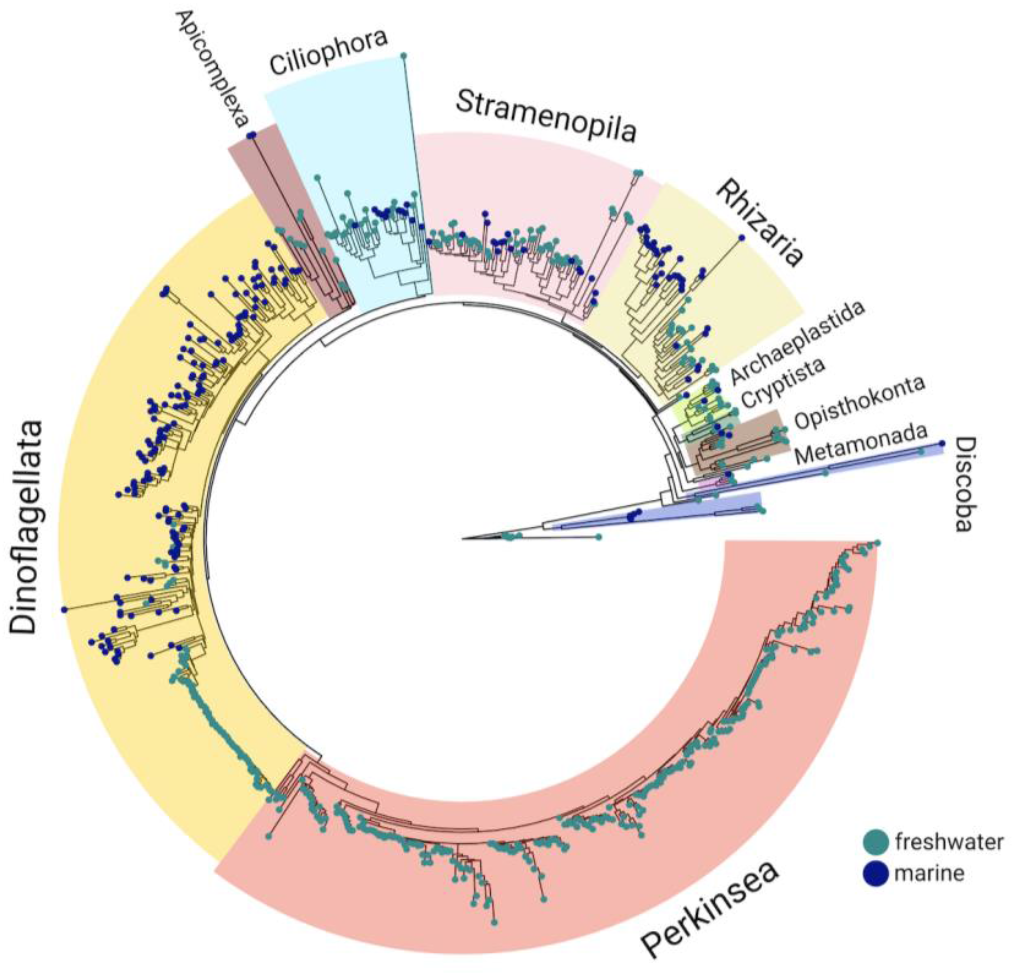
Phylogenetic tree of 721 novel 18S rDNA OTUs identified with Nanopore sequencing from both freshwater and marine environments. The tree was built with IQ-TREE using TIM2+F+R10 model.

Some new taxa, such as those belonging to Haptista, Ancyromonadida, Telonemia, Rhodophyta and Chrompodellids were only represented by a small number of sequences and did not form well defined groups, in contrast to the majority of sequences which formed monophyletic groups. Most of the novel OTUs belonged to the perkinseans and dinoflagellates (Alveolata), followed by gyristans, ciliophoreans, bigyrans, cercozoans and radiolarians. In freshwater the highest number of novel taxa belonged to classes Perkinsida (Perkinsea) with 254 OTUs, Dinophyceae (Dinoflagellata) with 73 OTUs, Chrysophyceae (Gyrista) with 19 OTUs and Bicoecea (Bigyra) with 11 OTUs. In marine environments they belonged mostly to Syndiniales (Dinoflagellata) with 106 OTUs, Dinophyceae (Dinoflagellata) with 46 OTUs, RAD-A (Radiolaria) with 13 OTUs and Spirotrichea (Ciliophora) with 10 OTUs.

## Discussion

### Generating high-quality OTUs through Nanopore sequencing and the BaNaNA pipeline

So far, one of the biggest challenges for metabarcoding based on Nanopore sequencing was the relatively high error rate of this technology. In the last years, continuous advances in the chemistry and base-calling tools have significantly reduced the error rate (Chang et al., 2024; Ni et al., 2023; Zhang et al., 2023). For instance, accuracy has increased from 97.6% with R9.4.1 to 99.1% with R10.4 (Sereika et al., 2022), and further improvements are expected in the future. Still, current pipelines for processing Nanopore amplicons must account for the high error rate and denoising protocols used for Illumina sequenced amplicons resulting in ASVs can’t be applied. Instead, we proposed to generate OTUs by clustering and creating consensus sequences by alignment and comparing each position inside the alignment followed by polishing steps (Fig. 1). Primarily designed for protists, the BaNaNa pipeline is currently the only pipeline for obtaining OTUs from Nanopore amplicons which was benchmarked on both mock community and environmental data. Unlike some approaches designed for bacterial 16S rRNA gene (Ammer-Herrmenau et al., 2021; Curry et al., 2022, EPI2ME-Oxford Nanopore Technologies) our pipeline creates OTUs independently of a reference database which is crucial in case of limited reference databases such as for protists. In fact only Natrix2 (Bludau et al., 2025) was so far benchmarked on eukaryotic sediment samples for protist and other eukaryotic meta-barcodes. BaNaNA is based on a similar approach to Natrix2, however it includes additional steps of quality check of extracted rDNA fragments (extracting_rrna.py) and it creates consensus using a custom script which aligns and compares each position in the alignment to eliminate potential sequencing errors (mafft_consensus.py). Additionally, the abundance calculations are based on counting sequences in clusters and not on the mapping of the final OTUs to the reads. Similarly to other existing approaches (Deep et al., 2023; Dubois et al., 2024; Egeter et al., 2022; Huggins et al., 2024; Lemoinne et al., 2024; Ohta et al., 2023; Rodríguez-Pérez et al., 2021; Schacksen et al., 2024) BaNaNA includes clustering steps which are crucial for metabarcoding of more complex communities and allow to obtain taxonomic units in a comparable manner between studies and to generate the abundance information.

### Evaluating Nanopore OTUs *versus* Illumina ASVs

Comparing two sequencing technologies and bioinformatic pipelines for obtaining representative sequences (ASVs and OTUs) is far from obvious as many parameters can potentially impact the outcome of the whole process. For example, amplification efficiency can decrease with longer amplicons (Arezi et al., 2003) and sequencing reproducibility is not perfect, even for a given technology, despite analysing all replicates using the exact same approach (Wen et al., 2017). With respect to the factors impacting the quantitative results, the varying number of copies of the 18S rRNA gene in different species is critical (Biard et al., 2017; W. Gong & Marchetti, 2019; Martin et al., 2022). High copy numbers in protist groups like ciliates (Biard et al., 2017; J. Gong et al., 2013; Martin et al., 2022) and dinoflagellates (Biard et al., 2017; W. Gong & Marchetti, 2019; Martin et al., 2022) promotes their amplification and overestimates their abundance, which resulted in increased relative abundances in the mock community (Fig. 2A). Another factor which can inflate estimated diversity is the intra-genomic rDNA gene diversity in the cell (Decelle et al., 2014; Sandin et al., 2022), where small differences between OTUs can be interpreted as different species, whereas these are just different variants within single species. Finally, rapid diversification of the lineage might result in high diversity as it is observed for perkinseans (Jobard et al., 2020). Indeed, in our data we obtained multiple ASVs and OTUs for one individual species, which can be the result of either small errors or detected variants. Our pipeline by using clustering instead of denoising to create MOTUs can remove some of the intra-species diversity, however a high number of closely related OTUs is still clearly visible on the phylogenetic tree (Fig. 5), especially within the Dinoflagellata and Perkinsea groups. The proper choice of primers also has a crucial impact on the final results, as there is no perfect pair of primers which would efficiently amplify all groups of protists (Latz et al., 2022; Vaulot et al., 2022). One group which is always underestimated by V4 Illumina primers are excavates (Choi & Park, 2020; Novák et al., 2024; Obiol et al., 2020) which have more mismatches in the place of primer annealing (Vaulot et al., 2022) and their longer V4 region (Geisen et al., 2015; Obiol et al., 2020) exceeds the Illumina read length capacity. Such bias towards excavates is clearly present in both mock community (Fig. 2A) and environmental community results obtained by Illumina sequencing (Fig. 4A,B) but was overcome by long-read Nanopore sequencing (Fig. 2A, Fig. 4A,B). Besides excavates, other groups like amoebozoans, rhizarians and Prymnesiophyceae are also underestimated in environmental samples due to the primers biases (Caron & Hu, 2019; Choi & Park, 2020; Obiol et al., 2020; Vaulot et al., 2022). It is possible that the low abundances of *Chlorella variabilis* and *Cryptomonas paramecium*, as well as the lack of detection of *Cryptomonas gyropyrenoidosa* in the Nanopore mock community dataset (Table 1) are due to low amplification efficiency for these particular species. This is especially likely given that the species had their representatives present in the reference database and both groups (Chlorophyta and Cryptophyta) were well represented in the environmental data (Fig. 4A).

Both Illumina and Nanopore sequencing revealed taxa that were not originally present in the mock community (Fig. 2A, Fig. S2), what we believe is mainly due to the artefacts in amplicon generation. This effect is particularly pronounced in the ASVs generated with DADA2, which yielded significantly more taxa than the OTUs derived from Nanopore data using the BaNaNA pipeline (Fig. S1, Fig. 2B). The high number of false positives associated with DADA2 was also observed in previous studies (e.g. Overgaard et al., 2024). This is also reflected in the almost exponential increase of taxa at lower taxonomic levels in the Illumina dataset, indicating a high level of background noise (Fig. S1), possibly leading to inflated diversity estimates in environmental samples (Fig. 3). In contrast, the Nanopore-based OTUs showed fewer false positives, suggesting a more accurate representation of actual protist diversity. Some non-mock taxa were shared across both platforms and may result from incidental presence of these taxa in cultures, e.g. Perkinsea, a known parasite of dinoflagellates, and *Cryptomonas paramecium* (=*Chilomonas paramecium*) (Brugerolle, 2002; Itoïz et al., 2022). Others could be caused by minor contamination during laboratory processing or by cross-contamination such as “tag-jumping” (Schnell et al., 2015). These are known sources of errors (Santoferrara, 2019), which however had no significant impact on the ecological interpretation of obtained results (Fig. 4A,B,D).

### Reconstructing ecological patterns from Nanopore and Illumina-based metabarcoding

Both sequencing technologies applied to environmental samples yielded similar taxonomic compositions (Fig. 4A,B) and successfully reconstructed ecological patterns (Fig. 4D). Such agreement has been previously demonstrated for long-read PacBio metabarcoding studies (Burki et al., 2021; Jamy et al., 2022) and Nanopore metabarcoding on protist communities (Bludau et al., 2025). Samples as expected were first differentiated based on the salinity of the environment dividing them into marine and freshwater ones (Fig. 4D). Furthermore, deep-sea samples were clearly separated from the ones collected at the surface and DCM layers of the water column. Sunlit communities were characterized with photosynthetic taxa such as chlorophytes and haptophytes whereas deep-sea samples were dominated by heterotrophic radiolarians and diplonemids (Fig. 4B). Survey of freshwater dimictic lakes from summer revealed an expected pattern with substantial presence of dinoflagellates and cryptophytes in oxygenated layers (Debroas et al., 2017; Karlicki et al., 2024) (Fig. 4A). The unclear distinction between the photic and aphotic zones could be due to the strong influence of sinking dead cells, as previously demonstrated (Karlicki et al., 2024). That is furthermore supported by higher relative abundance of parasitic Perkinsea in deeper fractions which has been noticed by other metabarcoding and microscopic surveys (Karlicki et al., 2024; Mangot et al., 2009).

### The importance of reference database in metabarcoding studies

The comprehensiveness of reference sequence database is crucial to properly describe protist diversity (Tragin et al., 2018). However, existing protist databases cover less diversity compared to the bacterial ones, thus making the taxonomic annotation less accurate. Additionally, it appears that there is a disproportion in the number of available protist reference sequences for marine and freshwater environments. Protists in marine ecosystems are better studied than freshwater ones with molecular methods and currently have much better representation of species in the databases. This disparity is well visible when looking at the percentage of identities of ASVs and OTUs to the closest reference (Fig. 4C) where generally lower identities for freshwater environment are caused by a missing close reference in the PR2 database. We also obtained twice as many novel full-length 18S rDNA OTUs from freshwater data than from marine (Fig. 5). Our novel full-length 18S rDNA OTUs represent many groups of protists (Fig. 5) clearly showing systematic gaps in the reference databases but also showing the potential to fill this gap and improve future classifications with long Nanopore meta-barcodes. The proper classification of long reads requires also databases with longer sequences; the primers tested here resulted in the full 18S rDNA, however, Nanopore sequencing allows to generate amplicons covering the whole rDNA operon, though such databases allowing appropriate classification of the full operon are still very limited (Krabberød et al., 2025; Tedersoo et al., 2024).

### Longer is better: the benefits of Nanopore amplicons

The shorter length of Illumina amplicons prevents accurate taxonomy resolution beyond the genus level, limiting evolutionary and fine scale ecological studies (Hugerth et al., 2014; Szoboszlay et al., 2023). Nanopore technology allows to sequence at once the whole 18S rRNA gene and more, providing access to more information and therefore a much better species-level resolution of taxonomic annotation (Bludau et al., 2025; Latz et al., 2022; Ohta et al., 2023; Pascoal et al., 2024; Petrone et al., 2023; Szoboszlay et al., 2023; Zhang et al., 2023). The decrease in annotation accuracy is particularly visible when comparing full-length 18S rDNA OTUs with V4-tags as the latter had fewer sequences assigned at the species level, which is especially visible for *Paramecium bursaria* in the mock community (Fig. 2B, Table 1). On the other hand, shorter sequences are more likely to match with 100% identity to a reference than longer sequences, as seen with V4-tags which generally exhibit higher identity percentages with the same reference database compared to full-length 18S rDNA OTUs (Fig. 4C) which may result with overestimation of taxa. In addition, unlike short ASVs, long OTUs are also suitable for phylogenetic reconstruction (Fig. 5) (Overgaard et al., 2024), what allows their taxonomy to be curated and potentially to discover new lineages (Lara et al., 2009).

Illumina amplicon sequencing is currently the gold standard for metabarcoding of protists communities (De Vargas et al., 2015; Ibarbalz et al., 2023; Karlicki et al., 2024; Mahé et al., 2017; Piredda et al., 2017) and vast amounts of data have been collected and deposited in public databases up till now. Still, the data are heterogeneous as they were often generated using different primers or covering different fragments of the 18S rRNA gene and thus preventing a proper datasets integration or comparison. Sequencing the whole 18S rRNA gene by Nanopore offers the possibility to use any primer pair to extract needed fragment of the gene and combine it with Illumina datasets, as demonstrated by our V4-tags analysis. Taxonomic assignment of the extracted fragment gives similar results, yet with a lower accuracy compared to the full-length 18S rDNA OTUs (Fig. 2, Fig. 4A, B). Therefore, we recommend to retain the taxonomic assignment from the whole full-length 18S rDNA to fully use the possibilities of long Nanopore OTUs before extracting any specific region of interest.

## Conclusions

This study shows the effectiveness of long-read Nanopore sequencing for protist biodiversity, ecology and evolution research, particularly in combination with our newly developed BaNaNA pipeline. Our results based on high-quality OTUs confirmed that Nanopore sequencing is a powerful and reliable tool for diversity studies, providing comparable results to Illumina while reducing the noise often associated with short-read technologies and improving taxonomic resolution. The ability to use those sequences for phylogenetic reconstructions and higher taxonomic resolution further highlights the advantages of longer amplicon data for ecological analyses.

We have also demonstrated that large amounts of Illumina-based metabarcoding data can be effectively combined with Nanopore meta-barcodes. The V4-tags extracted from Nanopore full-length 18S rDNA OTUs and Illumina V4-18S rDNA ASVs are highly comparable. This compatibility enables the integration of data generated with different sequencing technologies and primer pairs and facilitates long-term studies by incorporating existing data.

Finally, our analysis shows that marine samples with more complete reference databases benefit from the higher resolution of long-read sequencing. However, challenges in taxonomic annotation remain due to incomplete reference databases for freshwater and other environments, emphasizing the need to further improve protist reference databases to fully exploit long-reads metabarcoding potential.

## Data Availability Statement

The sequencing data have been deposited in the EMBL-EBI European Nucleotide Archive (ENA) under the Projects IDs PRJEB89945 and PRJEB76575. All the rest of supplementary materials can be found in Zenodo repository under DOI 10.5281/zenodo.15673958.

## Acknowledgements

Freshwater sampling was conducted using the facilities of the KUMAK Masurian Centre for Biodiversity and Education in Urwitałt, Faculty of Biology, University of Warsaw. We would like to thank all the MicroDivEr team members who helped us with the sampling. We acknowledge the MOOSE program (Mediterranean Ocean Observing System for the Environment) coordinated by CNRS-INSU and the Research Infrastructure ILICO (CNRS-IFREMER).

## Author Contributions

Conceptualization (AK), Methodology (MC, SR, MK), Software (MC, MK), Validation (MC), Formal analysis (MC), Investigation (AK, SR, MC, MK), Resources (AK), Data Curation (MC), Writing - Original draft (MC), Writing - Review and Editing (AK, SR, MC, MK, FN), Visualization (MC), Supervision (AK, FN), Project administration (AK, FN), Funding Acquisition (AK).

## Bibliography

Altschul, S. F., Gish, W., Miller, W., Myers, E. W., & Lipman, D. J. (1990). Basic local alignment search tool. Journal of Molecular Biology, 215(3), 403–410.

Amaral-Zettler, L. A., McCliment, E. A., Ducklow, H. W., & Huse, S. M. (2009). A Method for Studying Protistan Diversity Using Massively Parallel Sequencing of V9 Hypervariable Regions of Small-Subunit Ribosomal RNA Genes. PLoS ONE, 4(7), e6372. 10.1371/journal.pone.0006372

Ammer-Herrmenau, C., Pfisterer, N., Van Den Berg, T., Gavrilova, I., Amanzada, A., Singh, S. K., Khalil, A., Alili, R., Belda, E., Clement, K., Abd El Wahed, A., Gady, E. E., Haubrock, M., Beißbarth, T., Ellenrieder, V., & Neesse, A. (2021). Comprehensive Wet-Bench and Bioinformatics Workflow for Complex Microbiota Using Oxford Nanopore Technologies. mSystems, 6(4), 10.1128/msystems.00750-21. 10.1128/msystems.00750-21

Andrews, S. (2010). FastQC: A Quality Control Tool for High Throughput Sequence Data [Online] [Computer software]. Available online at http://www.bioinf ormatics.babraham.ac.uk/projects/fastqc/

Arezi, B., Xing, W., Sorge, J. A., & Hogrefe, H. H. (2003). Amplification efficiency of thermostable DNA polymerases. Analytical Biochemistry, 321(2), 226–235. 10.1016/S0003-2697(03)00465-2

Biard, T., Bigeard, E., Audic, S., Poulain, J., Gutierrez-Rodriguez, A., Pesant, S., Stemmann, L., & Not, F. (2017). Biogeography and diversity of Collodaria (Radiolaria) in the global ocean. The ISME Journal, 11(6), 1331–1344. 10.1038/ismej.2017.12

Bludau, D., Sieber, G., Shah, M., Deep, A., Boenigk, J., & Beisser, D. (2025). Breaking the Standard: Can Oxford Nanopore Technologies Sequencing Compete With Illumina in Protistan Amplicon Studies? Environmental DNA, 7(2), e70084. 10.1002/edn3.70084

Bolyen, E., Rideout, J. R., Dillon, M. R., Bokulich, N. A., Abnet, C. C., Al-Ghalith, G. A., Alexander, H., Alm, E. J., Arumugam, M., Asnicar, F., Bai, Y., Bisanz, J. E., Bittinger, K., Brejnrod, A., Brislawn, C. J., Brown, C. T., Callahan, B. J., Caraballo-Rodríguez, A. M., Chase, J., … Caporaso, J. G. (2019). Reproducible, interactive, scalable and extensible microbiome data science using QIIME 2. Nature Biotechnology, 37(8), 852–857. 10.1038/s41587-019-0209-9

Brugerolle, G. (2002). Cryptophagus subtilis: A new parasite of cryptophytes affiliated with the Perkinsozoa lineage. European Journal of Protistology, 37(4), 379–390. 10.1078/0932-4739-00837

Burki, F., Sandin, M. M., & Jamy, M. (2021). Diversity and ecology of protists revealed by metabarcoding. Current Biology, 31(19), R1267–R1280. 10.1016/j.cub.2021.07.066

Callahan, B. J., McMurdie, P. J., & Holmes, S. P. (2017). Exact sequence variants should replace operational taxonomic units in marker-gene data analysis. The ISME Journal, 11(12), 2639–2643. 10.1038/ismej.2017.119

Callahan, B. J., McMurdie, P. J., Rosen, M. J., Han, A. W., Johnson, A. J. A., & Holmes, S. P. (2016). DADA2: High-resolution sample inference from Illumina amplicon data. Nature Methods, 13(7), 581–583. 10.1038/nmeth.3869

Callahan, B. J., Wong, J., Heiner, C., Oh, S., Theriot, C. M., Gulati, A. S., McGill, S. K., & Dougherty, M. K. (2019). High-throughput amplicon sequencing of the full-length 16S rRNA gene with single-nucleotide resolution. Nucleic Acids Research, 47(18), e103–e103. 10.1093/nar/gkz569

Capella-Gutiérrez, S., Silla-Martínez, J. M., & Gabaldón, T. (2009). trimAl: A tool for automated alignment trimming in large-scale phylogenetic analyses. Bioinformatics, 25(15), 1972–1973.

Caron, D. A., & Hu, S. K. (2019). Are We Overestimating Protistan Diversity in Nature? Trends in Microbiology, 27(3), 197–205. 10.1016/j.tim.2018.10.009

Chang, J. J. M., Ip, Y. C. A., Neo, W. L., Mowe, M. A. D., Jaafar, Z., & Huang, D. (2024). Primed and ready: Nanopore metabarcoding can now recover highly accurate consensus barcodes that are generally indel-free. BMC Genomics, 25(1), 842. 10.1186/s12864-024-10767-4

Choi, J., & Park, J. S. (2020). Comparative analyses of the V4 and V9 regions of 18S rDNA for the extant eukaryotic community using the Illumina platform. Scientific Reports, 10(1), 6519. 10.1038/s41598-020-63561-z

Cock, P. J. A., Antao, T., Chang, J. T., Chapman, B. A., Cox, C. J., Dalke, A., Friedberg, I., Hamelryck, T., Kauff, F., Wilczynski, B., & De Hoon, M. J. L. (2009). Biopython: Freely available Python tools for computational molecular biology and bioinformatics. Bioinformatics, 25(11), 1422–1423. 10.1093/bioinformatics/btp163

Coppola, L., Raimbault, P., Mortier, L., & Testor, P. (2019). Monitoring the Environment in the Northwestern Mediterranean Sea. Eos, 100. 10.1029/2019EO125951

Curry, K. D., Wang, Q., Nute, M. G., Tyshaieva, A., Reeves, E., Soriano, S., Wu, Q., Graeber, E., Finzer, P., Mendling, W., Savidge, T., Villapol, S., Dilthey, A., & Treangen, T. J. (2022). Emu: Species-level microbial community profiling of full-length 16S rRNA Oxford Nanopore sequencing data. Nature Methods, 19(7), 845–853. 10.1038/s41592-022-01520-4

De Coster, W., & Rademakers, R. (2023). NanoPack2: Population-scale evaluation of long-read sequencing data. Bioinformatics, 39(5), btad311. 10.1093/bioinformatics/btad311

De Vargas, C., Audic, S., Henry, N., Decelle, J., Mahé, F., Logares, R., Lara, E., Berney, C., Le Bescot, N., Probert, I., Carmichael, M., Poulain, J., Romac, S., Colin, S., Aury, J.-M., Bittner, L., Chaffron, S., Dunthorn, M., Engelen, S., … Velayoudon, D. (2015). Eukaryotic plankton diversity in the sunlit ocean. Science, 348(6237), 1261605. 10.1126/science.1261605

Debroas, D., Domaizon, I., Humbert, J.-F., Jardillier, L., Lepère, C., Oudart, A., & Taïb, N. (2017). Overview of freshwater microbial eukaryotes diversity: A first analysis of publicly available metabarcoding data. FEMS Microbiology Ecology, 93(4). 10.1093/femsec/fix023

Decelle, J., Romac, S., Sasaki, E., Not, F., & Mahé, F. (2014). Intracellular Diversity of the V4 and V9 Regions of the 18S rRNA in Marine Protists (Radiolarians) Assessed by High-Throughput Sequencing. PLoS ONE, 9(8), e104297. 10.1371/journal.pone.0104297

Deep, A., Bludau, D., Welzel, M., Clemens, S., Heider, D., Boenigk, J., & Beisser, D. (2023). Natrix2 – Improved amplicon workflow with novel Oxford Nanopore Technologies support and enhancements in clustering, classification and taxonomic databases. Metabarcoding and Metagenomics, 7, e109389. 10.3897/mbmg.7.109389

Dubois, B., Delitte, M., Lengrand, S., Bragard, C., Legrève, A., & Debode, F. (2024). PRONAME: A user-friendly pipeline to process long-read nanopore metabarcoding data by generating high-quality consensus sequences. Frontiers in Bioinformatics, 4, 1483255. 10.3389/fbinf.2024.1483255

Edgar, R. C. (2017). Accuracy of microbial community diversity estimated by closed- and open-reference OTUs. PeerJ, 5, e3889. 10.7717/peerj.3889

Edgcomb, V., Orsi, W., Bunge, J., Jeon, S., Christen, R., Leslin, C., Holder, M., Taylor, G. T., Suarez, P., Varela, R., & Epstein, S. (2011). Protistan microbial observatory in the Cariaco Basin, Caribbean. I. Pyrosequencing vs Sanger insights into species richness. The ISME Journal, 5(8), 1344–1356. 10.1038/ismej.2011.6

Egeter, B., Veríssimo, J., Lopes-Lima, M., Chaves, C., Pinto, J., Riccardi, N., Beja, P., & Fonseca, N. A. (2022). Speeding up the detection of invasive bivalve species using environmental DNA: A Nanopore and Illumina sequencing comparison. Molecular Ecology Resources, 22(6), 2232–2247. 10.1111/1755-0998.13610

Gaonkar, C. C., & Campbell, L. (2024). A full-length 18S ribosomal DNA metabarcoding approach for determining protist community diversity using Nanopore sequencing. Ecology and Evolution, 14(4), e11232. 10.1002/ece3.11232

Geisen, S., Laros, I., Vizcaíno, A., Bonkowski, M., & De Groot, G. A. (2015). Not all are free-living: High-throughput DNA metabarcoding reveals a diverse community of protists parasitizing soil metazoa. Molecular Ecology, 24(17), 4556–4569. 10.1111/mec.13238

Gong, J., Dong, J., Liu, X., & Massana, R. (2013). Extremely High Copy Numbers and Polymorphisms of the rDNA Operon Estimated from Single Cell Analysis of Oligotrich and Peritrich Ciliates. Protist, 164(3), 369–379. 10.1016/j.protis.2012.11.006

Gong, W., & Marchetti, A. (2019). Estimation of 18S Gene Copy Number in Marine Eukaryotic Plankton Using a Next-Generation Sequencing Approach. Frontiers in Marine Science, 6, 219. 10.3389/fmars.2019.00219

Guillou, L., Bachar, D., Audic, S., Bass, D., Berney, C., Bittner, L., Boutte, C., Burgaud, G., De Vargas, C., Decelle, J., Del Campo, J., Dolan, J. R., Dunthorn, M., Edvardsen, B., Holzmann, M., Kooistra, W. H. C. F., Lara, E., Le Bescot, N., Logares, R., … Christen, R. (2012). The Protist Ribosomal Reference database (PR2): A catalog of unicellular eukaryote Small Sub-Unit rRNA sequences with curated taxonomy. Nucleic Acids Research, 41(D1), D597–D604. 10.1093/nar/gks1160

Hooper, C., Ward, G. M., Foster, R., Skujina, I., Ironside, J. E., Berney, C., & Bass, D. (2023). Long amplicons as a tool to identify variable regions of ribosomal RNA for improved taxonomic resolution and diagnostic assay design in microeukaryotes: Using ascetosporea as a case study. Frontiers in Ecology and Evolution, 11, 1266151. 10.3389/fevo.2023.1266151

Hugerth, L. W., Muller, E. E. L., Hu, Y. O. O., Lebrun, L. A. M., Roume, H., Lundin, D., Wilmes, P., & Andersson, A. F. (2014). Systematic Design of 18S rRNA Gene Primers for Determining Eukaryotic Diversity in Microbial Consortia. PLoS ONE, 9(4), e95567. 10.1371/journal.pone.0095567

Huggins, L. G., Colella, V., Young, N. D., & Traub, R. J. (2024). Metabarcoding using nanopore long-read sequencing for the unbiased characterization of apicomplexan haemoparasites. Molecular Ecology Resources, 24(2), e13878. 10.1111/1755-0998.13878

Ibarbalz, F. M., Henry, N., Mahé, F., Ardyna, M., Zingone, A., Scalco, E., Lovejoy, C., Lombard, F., Jaillon, O., Iudicone, D., Malviya, S., Tara Oceans Coordinators, Sullivan, M. B., Chaffron, S., Karsenti, E., Babin, M., Boss, E., Wincker, P., Zinger, L., … Karp-Boss, L. (2023). Pan-Arctic plankton community structure and its global connectivity. Elementa: Science of the Anthropocene, 11(1), 00060. 10.1525/elementa.2022.00060

Itoïz, S., Metz, S., Derelle, E., Reñé, A., Garcés, E., Bass, D., Soudant, P., & Chambouvet, A. (2022). Emerging Parasitic Protists: The Case of Perkinsea. Frontiers in Microbiology, 12, 735815. 10.3389/fmicb.2021.735815

Jamy, M., Biwer, C., Vaulot, D., Obiol, A., Jing, H., Peura, S., Massana, R., & Burki, F. (2022). Global patterns and rates of habitat transitions across the eukaryotic tree of life. Nature Ecology & Evolution, 6(10), 1458–1470. 10.1038/s41559-022-01838-4

Jamy, M., Foster, R., Barbera, P., Czech, L., Kozlov, A., Stamatakis, A., Bending, G., Hilton, S., Bass, D., & Burki, F. (2020). Long-read metabarcoding of the eukaryotic rDNA operon to phylogenetically and taxonomically resolve environmental diversity. Molecular Ecology Resources, 20(2), 429–443. 10.1111/1755-0998.13117

Jobard, M., Wawrzyniak, I., Bronner, G., Marie, D., Vellet, A., Sime-Ngando, T., Debroas, D., & Lepère, C. (2020). Freshwater Perkinsea: Diversity, ecology and genomic information. Journal of Plankton Research, 42(1), 3–17. 10.1093/plankt/fbz068

Karlicki, M., Bednarska, A., Hałakuc, P., Maciszewski, K., & Karnkowska, A. (2024). Spatio-temporal changes of small protist and free-living bacterial communities in a temperate dimictic lake: Insights from metabarcoding and machine learning. FEMS Microbiology Ecology, 100(8), fiae104. 10.1093/femsec/fiae104

Katoh, K., & Standley, D. M. (2013). MAFFT Multiple Sequence Alignment Software Version 7: Improvements in Performance and Usability. Molecular Biology and Evolution, 30(4), 772–780. 10.1093/molbev/mst010

Krabberød, A. K., Stokke, E., Thoen, E., Skrede, I., & Kauserud, H. (2025). The Ribosomal Operon Database: A Full-Length RDNA Operon Database Derived From Genome Assemblies. Molecular Ecology Resources, 25(1), e14031. 10.1111/1755-0998.14031

Lara, E., Moreira, D., Vereshchaka, A., & López-García, P. (2009). Pan-oceanic distribution of new highly diverse clades of deep-sea diplonemids. Environmental Microbiology, 11(1), 47–55. 10.1111/j.1462-2920.2008.01737.x

Latz, M. A. C., Grujcic, V., Brugel, S., Lycken, J., John, U., Karlson, B., Andersson, A., & Andersson, A. F. (2022). Short- and long-read metabarcoding of the eukaryotic rRNA operon: Evaluation of primers and comparison to shotgun metagenomics sequencing. Molecular Ecology Resources, 22(6), 2304–2318. 10.1111/1755-0998.13623

Lemoinne, A., Dirberg, G., Georges, M., & Robinet, T. (2024). Evaluation of a Nanopore Sequencing Strategy on Bacterial Communities From Marine Sediments. Environmental DNA, 6(5), e70009. 10.1002/edn3.70009

Li, H. (2018). Minimap2: Pairwise alignment for nucleotide sequences. Bioinformatics, 34(18), 3094–3100. 10.1093/bioinformatics/bty191

Mahé, F., De Vargas, C., Bass, D., Czech, L., Stamatakis, A., Lara, E., Singer, D., Mayor, J., Bunge, J., Sernaker, S., Siemensmeyer, T., Trautmann, I., Romac, S., Berney, C., Kozlov, A., Mitchell, E. A. D., Seppey, C. V. W., Egge, E., Lentendu, G., … Dunthorn, M. (2017). Parasites dominate hyperdiverse soil protist communities in Neotropical rainforests. Nature Ecology & Evolution, 1(4), 0091. 10.1038/s41559-017-0091

Mangot, J.-F., Lepère, C., Bouvier, C., Debroas, D., & Domaizon, I. (2009). Community Structure and Dynamics of Small Eukaryotes Targeted by New Oligonucleotide Probes: New Insight into the Lacustrine Microbial Food Web. Applied and Environmental Microbiology, 75(19), 6373–6381. 10.1128/AEM.00607-09

Martin, J. L., Santi, I., Pitta, P., John, U., & Gypens, N. (2022). Towards quantitative metabarcoding of eukaryotic plankton: An approach to improve 18S rRNA gene copy number bias. Metabarcoding and Metagenomics, 6, e85794. 10.3897/mbmg.6.85794

McMurdie, P. J., & Holmes, S. (2013). phyloseq: An R package for reproducible interactive analysis and graphics of microbiome census data. PLoS ONE, 8(4), e61217.

Medlin, L., Elwood, H. J., Stickel, S., & Sogin, M. L. (1988). The characterization of enzymatically amplified eukaryotic 16S-like rRNA-coding regions. Gene, 71(2), 491–499.

Minh, B. Q., Schmidt, H. A., Chernomor, O., Schrempf, D., Woodhams, M. D., Von Haeseler, A., & Lanfear, R. (2020). IQ-TREE 2: New models and efficient methods for phylogenetic inference in the genomic era. Molecular Biology and Evolution, 37(5), 1530–1534.

Mölder, F., Jablonski, K. P., Letcher, B., Hall, M. B., Tomkins-Tinch, C. H., Sochat, V., Forster, J., Lee, S., Twardziok, S. O., Kanitz, A., & others. (2021). Sustainable data analysis with Snakemake. F1000Research, 10.

Ni, Y., Liu, X., Simeneh, Z. M., Yang, M., & Li, R. (2023). Benchmarking of Nanopore R10.4 and R9.4.1 flow cells in single-cell whole-genome amplification and whole-genome shotgun sequencing. Computational and Structural Biotechnology Journal, 21, 2352–2364. 10.1016/j.csbj.2023.03.038

Novák, J., Treitli, S. C., Füssy, Z., Záhonová, K., Hamplová, B., Hrdá, Š., & Hampl, V. (2024). V9 Hypervariable Region Metabarcoding Primers for Euglenozoa and Metamonada. Environmental DNA, 6(5), e70022. 10.1002/edn3.70022

Obiol, A., Giner, C. R., Sánchez, P., Duarte, C. M., Acinas, S. G., & Massana, R. (2020). A metagenomic assessment of microbial eukaryotic diversity in the global ocean. Molecular Ecology Resources, 20(3), 718–731. 10.1111/1755-0998.13147

Ohta, A., Nishi, K., Hirota, K., & Matsuo, Y. (2023). Using nanopore sequencing to identify fungi from clinical samples with high phylogenetic resolution. Scientific Reports, 13(1), 9785. 10.1038/s41598-023-37016-0

Oksanen, J., Simpson, G. L., Blanchet, F. G., Kindt, R., Legendre, P., Minchin, P. R., O’Hara, R. B., Solymos, P., Stevens, M. H. H., Szoecs, E., Wagner, H., Barbour, M., Bedward, M., Bolker, B., Borcard, D., Carvalho, G., Chirico, M., Caceres, M. D., Durand, S., … Weedon, J. (2022). vegan: Community Ecology Package. https://CRAN.R-project.org/package=vegan

Olivier, S. A., Bull, M. K., Strube, M. L., Murphy, R., Ross, T., Bowman, J. P., & Chapman, B. (2023). Long-read MinION™ sequencing of 16S and 16S-ITS-23S rRNA genes provides species-level resolution of Lactobacillaceae in mixed communities. Frontiers in Microbiology, 14, 1290756. 10.3389/fmicb.2023.1290756

Overgaard, C. K., Jamy, M., Radutoiu, S., Burki, F., & Dueholm, M. K. D. (2024). Benchmarking long-read sequencing strategies for obtaining ASV -resolved RRNA operons from environmental microeukaryotes. Molecular Ecology Resources, 24(7), e13991. 10.1111/1755-0998.13991

Pagès, H., Aboyoun, P., Gentleman, R., & DebRoy, S. (2021). Biostrings: Efficient manipulation of biological strings. https://bioconductor.org/packages/Biostrings

Pascoal, F., Duarte, P., Assmy, P., Costa, R., & Magalhães, C. (2024). Full-length 16S rRNA gene sequencing combined with adequate database selection improves the description of Arctic marine prokaryotic communities. Annals of Microbiology, 74(1), 29. 10.1186/s13213-024-01767-6

Petrone, J. R., Rios Glusberger, P., George, C. D., Milletich, P. L., Ahrens, A. P., Roesch, L. F. W., & Triplett, E. W. (2023). RESCUE: A validated Nanopore pipeline to classify bacteria through long-read, 16S-ITS-23S rRNA sequencing. Frontiers in Microbiology, 14, 1201064. 10.3389/fmicb.2023.1201064

Piredda, R., Tomasino, M. P., D’Erchia, A. M., Manzari, C., Pesole, G., Montresor, M., Kooistra, W. H. C. F., Sarno, D., & Zingone, A. (2017). Diversity and temporal patterns of planktonic protist assemblages at a Mediterranean Long Term Ecological Research site. FEMS Microbiology Ecology, 93(1), fiw200. 10.1093/femsec/fiw200

Rodríguez-Pérez, H., Ciuffreda, L., & Flores, C. (2021). NanoCLUST: A species-level analysis of 16S rRNA nanopore sequencing data. Bioinformatics, 37(11), 1600–1601. 10.1093/bioinformatics/btaa900

Rognes, T., Flouri, T., Nichols, B., Quince, C., & Mahé, F. (2016). VSEARCH: A versatile open source tool for metagenomics. PeerJ, 4, e2584. 10.7717/peerj.2584

RStudio Team. (2020). RStudio: Integrated Development Environment for R [Computer software]. http://www.rstudio.com/

Sandin, M. M., Romac, S., & Not, F. (2022). Intra-genomic RRNA gene variability of Nassellaria and Spumellaria (Rhizaria, Radiolaria) assessed by Sanger, MINION and Illumina sequencing. Environmental Microbiology, 24(7), 2979–2993. 10.1111/1462-2920.16081

Santoferrara, L. F. (2019). Current practice in plankton metabarcoding: Optimization and error management. Journal of Plankton Research, 41(5), 571–582. 10.1093/plankt/fbz041

Santos, A., Van Aerle, R., Barrientos, L., & Martinez-Urtaza, J. (2020). Computational methods for 16S metabarcoding studies using Nanopore sequencing data. Computational and Structural Biotechnology Journal, 18, 296–305. 10.1016/j.csbj.2020.01.005

Schacksen, P. S., Østergaard, S. K., Eskildsen, M. H., & Nielsen, J. L. (2024). Complete pipeline for Oxford Nanopore Technology amplicon sequencing (ONT - AMPSEQ): From pre-processing to creating an operational taxonomic unit table. FEBS Open Bio, 14(11), 1779–1787. 10.1002/2211-5463.13868

Schnell, I. B., Bohmann, K., & Gilbert, M. T. P. (2015). Tag jumps illuminated – reducing sequence-to-sample misidentifications in metabarcoding studies. Molecular Ecology Resources, 15(6), 1289–1303. 10.1111/1755-0998.12402

Scholin, C. A., Herzog, M., Sogin, M., & Anderson, D. M. (1994). Identification of group-and strain-specific genetic markers for globally distributed Alexandrium (Dinophyceae). Ii. Sequence analysis of a fragment of the LSU rRNA gene 1. Journal of Phycology, 30(6), 999–1011.

Sereika, M., Kirkegaard, R. H., Karst, S. M., Michaelsen, T. Y., Sørensen, E. A., Wollenberg, R. D., & Albertsen, M. (2022). Oxford Nanopore R10.4 long-read sequencing enables the generation of near-finished bacterial genomes from pure cultures and metagenomes without short-read or reference polishing. Nature Methods, 19(7), 823–826. 10.1038/s41592-022-01539-7

Stoeck, T., Bass, D., Nebel, M., Christen, R., Jones, M. D. M., Breiner, H., & Richards, T. A. (2010). Multiple marker parallel tag environmental DNA sequencing reveals a highly complex eukaryotic community in marine anoxic water. Molecular Ecology, 19(1), 21–31. 10.1111/j.1365-294X.2009.04480.x

Szoboszlay, M., Schramm, L., Pinzauti, D., Scerri, J., Sandionigi, A., & Biazzo, M. (2023). Nanopore Is Preferable over Illumina for 16S Amplicon Sequencing of the Gut Microbiota When Species-Level Taxonomic Classification, Accurate Estimation of Richness, or Focus on Rare Taxa Is Required. Microorganisms, 11(3), 804. 10.3390/microorganisms11030804

Tedersoo, L., Hosseyni Moghaddam, M. S., Mikryukov, V., Hakimzadeh, A., Bahram, M., Nilsson, R. H., Yatsiuk, I., Geisen, S., Schwelm, A., Piwosz, K., Prous, M., Sildever, S., Chmolowska, D., Rueckert, S., Skaloud, P., Laas, P., Tines, M., Jung, J.-H., Choi, J. H., … Anslan, S. (2024). EUKARYOME: The rRNA gene reference database for identification of all eukaryotes. Database, 2024, baae043. 10.1093/database/baae043

Tragin, M., Zingone, A., & Vaulot, D. (2018). Comparison of coastal phytoplankton composition estimated from the V4 and V9 regions of the 18S rRNA gene with a focus on photosynthetic groups and especially Chlorophyta. Environmental Microbiology, 20(2), 506–520. 10.1111/1462-2920.13952

Vaser, R., Sović, I., Nagarajan, N., & Šikić, M. (2017). Fast and accurate de novo genome assembly from long uncorrected reads. Genome Research, 27(5), 737–746. 10.1101/gr.214270.116

Vaulot, D., Geisen, S., Mahé, F., & Bass, D. (2022). pr2-primers: An 18S rRNA primer database for protists. Molecular Ecology Resources, 22(1), 168–179. 10.1111/1755-0998.13465

Wang, L.-G., Lam, T. T.-Y., Xu, S., Dai, Z., Zhou, L., Feng, T., Guo, P., Dunn, C. W., Jones, B. R., Bradley, T., Zhu, H., Guan, Y., Jiang, Y., & Yu, G. (2020). treeio: An R package for phylogenetic tree input and output with richly annotated and associated data. Molecular Biology and Evolution, 37(2), 599–603. 10.1093/molbev/msz240

Wen, C., Wu, L., Qin, Y., Van Nostrand, J. D., Ning, D., Sun, B., Xue, K., Liu, F., Deng, Y., Liang, Y., & Zhou, J. (2017). Evaluation of the reproducibility of amplicon sequencing with Illumina MiSeq platform. PLOS ONE, 12(4), e0176716. 10.1371/journal.pone.0176716

Wheeler, D. L., Barrett, T., Benson, D. A., Bryant, S. H., Canese, K., Chetvernin, V., Church, D. M., DiCuccio, M., Edgar, R., Federhen, S., & others. (2007). Database resources of the national center for biotechnology information. Nucleic Acids Research, 36(Suppl_1), D13–D21.

Wickham, H. (2007). Reshaping Data with the reshape Package. Journal of Statistical Software, 21(12), 1–20.

Wickham, H. (2016). ggplot2: Elegant Graphics for Data Analysis. Springer-Verlag New York. https://ggplot2.tidyverse.org

Wickham, H., Averick, M., Bryan, J., Chang, W., McGowan, L. D., François, R., Grolemund, G., Hayes, A., Henry, L., Hester, J., Kuhn, M., Pedersen, T. L., Miller, E., Bache, S. M., Müller, K., Ooms, J., Robinson, D., Seidel, D. P., Spinu, V., … Yutani, H. (2019). Welcome to the tidyverse. Journal of Open Source Software, 4(43), 1686. 10.21105/joss.01686

Wickham, H., & Bryan, J. (2019). readxl: Read Excel Files. https://CRAN.R-project.org/package=readxl

Wickham, H., François, R., Henry, L., & Müller, K. (2022). dplyr: A Grammar of Data Manipulation. https://CRAN.R-project.org/package=dplyr

Yeh, Y.-C., & Fuhrman, J. A. (2022). Contrasting diversity patterns of prokaryotes and protists over time and depth at the San-Pedro Ocean Time series. ISME Communications, 2(1), 36. 10.1038/s43705-022-00121-8

Yu, G., Smith, D. K., Zhu, H., Guan, Y., & Lam, T. T.-Y. (2017). ggtree: An R package for visualization and annotation of phylogenetic trees with their covariates and other associated data. Methods in Ecology and Evolution, 8(1), 28–36.

Zhang, T., Li, H., Ma, S., Cao, J., Liao, H., Huang, Q., & Chen, W. (2023). The newest Oxford Nanopore R10.4.1 full-length 16S rRNA sequencing enables the accurate resolution of species-level microbial community profiling. Applied and Environmental Microbiology, 89(10), e00605–23. 10.1128/aem.00605-23

