## Supplementary Methods and Figures for "From short to long reads: enhanced protist diversity profiling via Nanopore metabarcoding"

### Detailed materials and methods

##### 1. Amplification of 18S rRNA gene fragment of *Cryptomonas gyropyrenoidosa* SAG 25.80 strain

To amplify 18S rRNA gene from *Cryptomonas gyropyrenoidosa* SAG 25.80 strain we used SA (5' AACCTGGTTGATCCTGCCAGT 3') and EukB (5' TGATCCTTCTGCAGGTTCACCTAC 3') (**Medlin et al., 1988**) primers (each at the final concentration of 0.2 µM). The 25 µl PCR mixture contained also 1µl of DNA template, 0.8 mM of mix of dNTPs, 1X of Phusion HF Buffer and 0.25 µl of Phusion High-Fidelity DNA polymerase (ThermoFisher). The program included: initial denaturation at 98°C for 30 s, 27 cycles of 10 s at 98°C, 30 s at 67°C, 1 m at 72°C and final elongation at 72°C for 5 m.

##### 2. Amplification of V4-18S rRNA gene fragment from freshwater samples and mock community

The amplification of the V4 region of 18S rRNA gene was performed in three replicates per sample in a 25 µl PCR mixture containing TAReuk454FWD1 (5' CCAGCASCYGCGGTAATTCC 3') and TAReukREV3 (5' ACTTTCGTTCTTGATYRA 3') (**Stoeck et al., 2010**) primers (each at final concentration of 0.2 µM), 1µl (5 ng/µl) of DNA template, 0.8 mM of mix of dNTPs, 1X of Phusion GC Buffer and 0.25 µl of Phusion High-Fidelity DNA polymerase (ThermoFisher). The program contined following steps: initial denaturation at 98°C for 30 s, 12 cycles of 10 s at 98°C, 30 s at 53°C, 30 s at 72°C, 18 cycles of 10 s at 98°C, 30 s at 48°C, 30 s at 72°C and final elongation at 72°C for 10 m.

##### 3. Amplification of 18S – D2 28S rRNA gene from freshwater and marine samples and mock community

We used SA (5' **TTTCTGTTGGTGCTGATATTGC**AACCTGGTTGATCCTGCCAGT 3') (**Medlin et al., 1988**) and D2C-R (5' **ACTTGCCTGTCGCTCTATCTTC**CCTTGGTCCGTGTTTCAAGA 3') (**Scholin et al., 1994**) primers to amplify a fragment from the beginning of 18S to D2 fragment of 28S rDNA. The primers are extended by an adapter sequence (**bold**), which is required by the PCR Barcoding Expansion 1-12 (EXP-PBC001) kit. The 25 µl PCR mixture contained primers in final concentration of 0.2 µM, 1µl (5 ng/µl) of DNA template, 1.6 mM of mix of dNTPs, 0.25 µl of Phusion High-Fidelity DNA polymerase (ThermoFisher, Finnzymes) and 1X of Phusion HF Buffer or Phusion GC Buffer respectively for freshwater and marine samples. The program contined following steps: initial denaturation at 98°C for 30 s, 27 cycles of 10 s at 98°C, 30 s at 59°C for freshwater samples or 55°C for marine samples, 2 m at 72°C and final elongation at 72°C for 10 m. The reaction was performed in three replicates per sample.

### References

Medlin, L., Elwood, H. J., Stickel, S., & Sogin, M. L. (1988). The characterization of enzymatically amplified eukaryotic 16S-like rRNA-coding regions. *Gene*, *71*(2), 491–499.

Scholin, C. A., Herzog, M., Sogin, M., & Anderson, D. M. (1994). Identification of group-and strain-specific genetic markers for globally distributed Alexandrium (Dinophyceae). Ii. Sequence analysis of a fragment of the LSU rRNA gene 1. *Journal of Phycology*, *30*(6), 999–1011.

Stoeck, T., Bass, D., Nebel, M., Christen, R., Jones, M. D. M., Breiner, H., & Richards, T. A. (2010). Multiple marker parallel tag environmental DNA sequencing reveals a highly complex eukaryotic community in marine anoxic water. *Molecular Ecology*, *19*(s1), 21–31. https://doi.org/10.1111/j.1365-294X.2009.04480.x

#### Supplementary figures

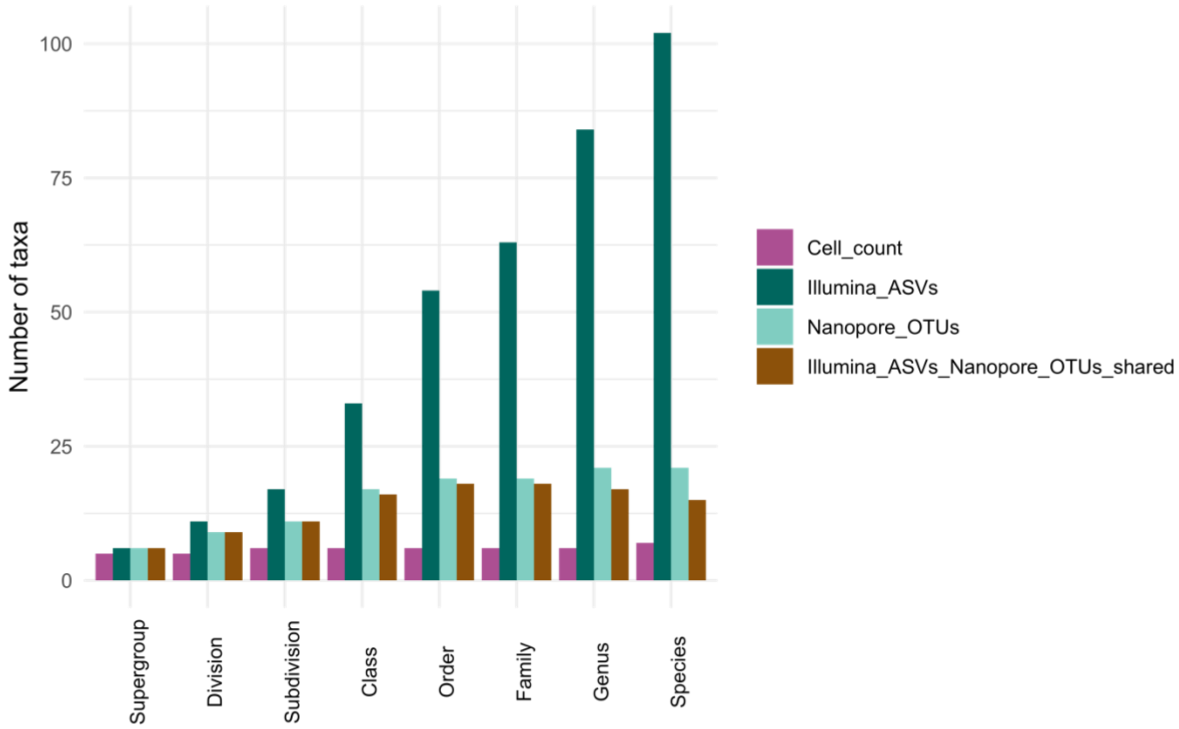


**Supplementary Figure 1** Number of taxa detected at different taxonomic ranks in the mock community determined by cell count and using Illumina and Nanopore sequencing. The figure shows the number of taxa identified separately in the Illumina and Nanopore datasets, as well as the number of taxa shared by both methods.

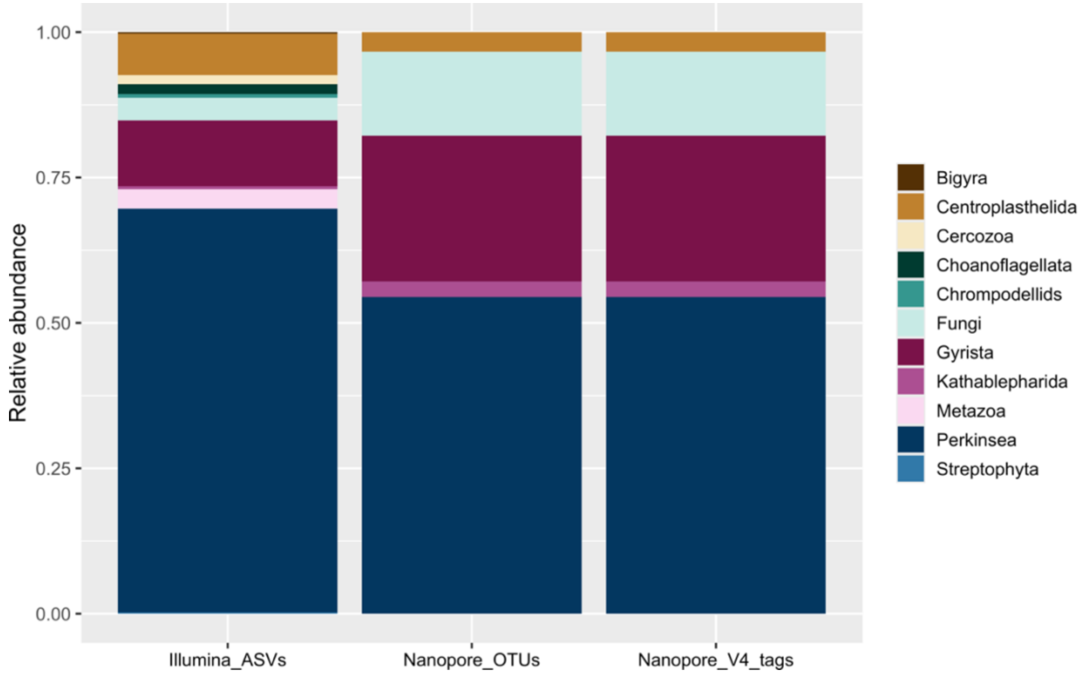


**Supplementary Figure 2** Relative abundance of species in the mock community classified as “Other” at the subdivision level, as determined by all sequencing approaches.

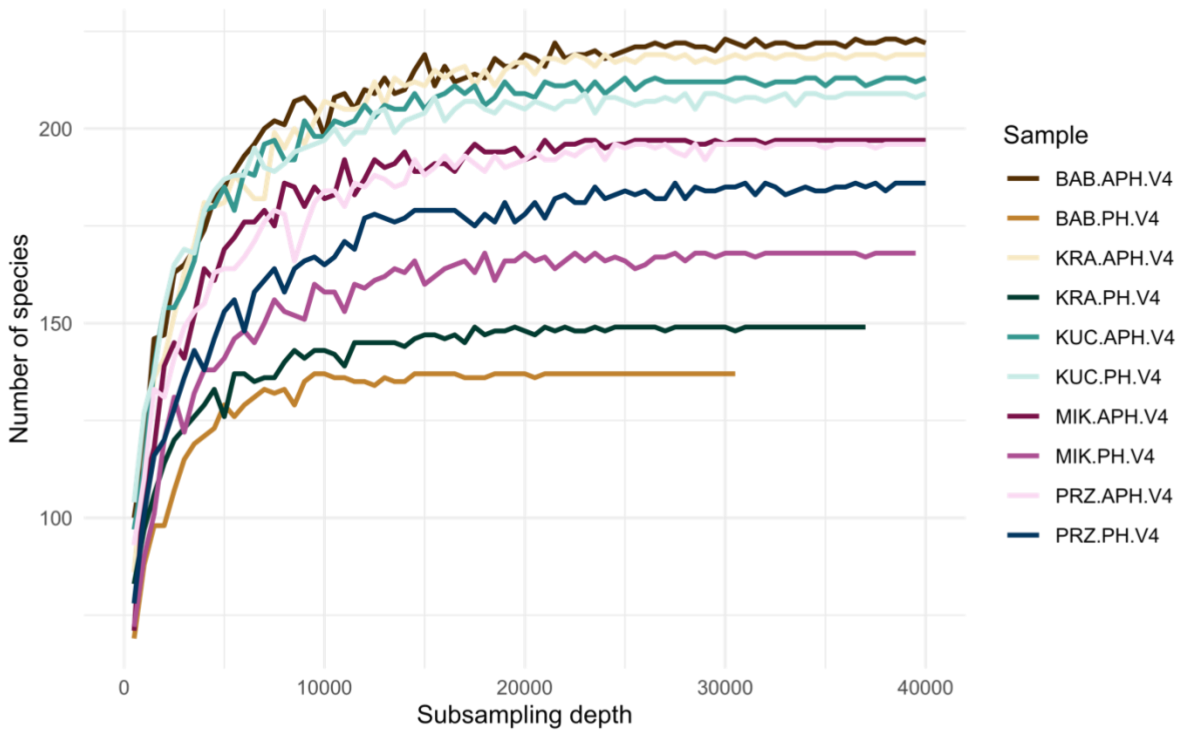


**Supplementary Figure 3** Rarefaction curves showing the number of observed species *per* freshwater sample sequenced with Illumina technology.

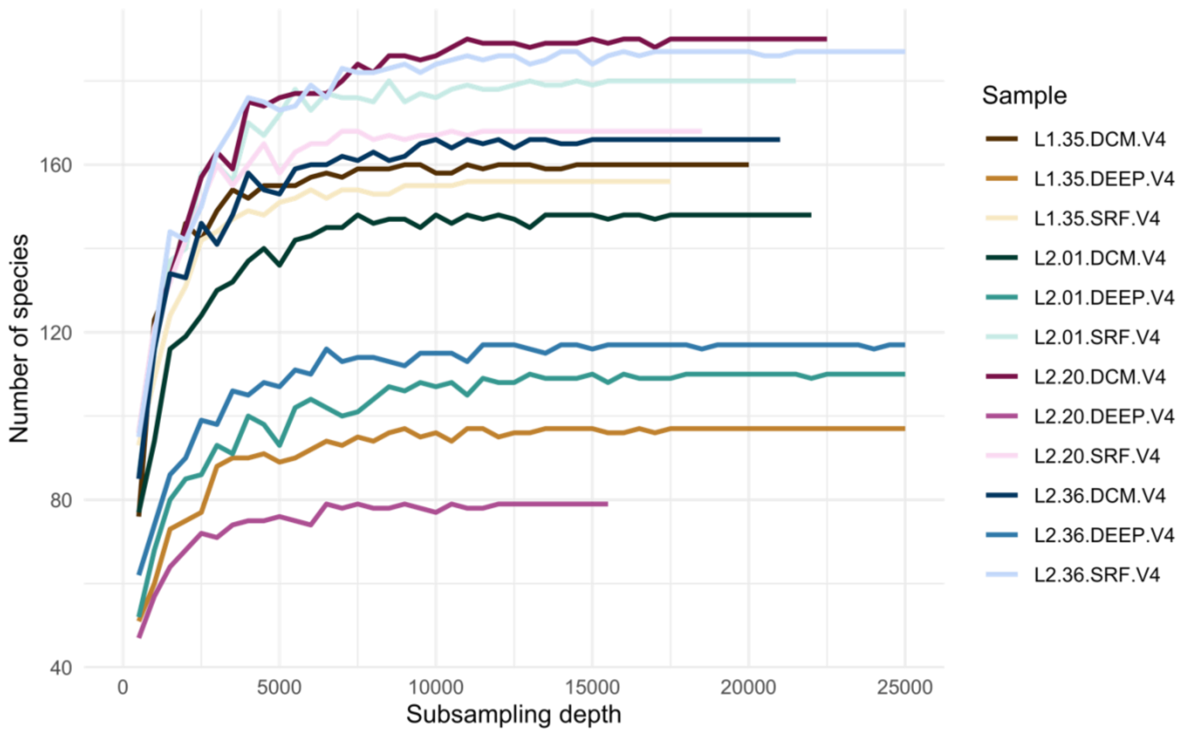


**Supplementary Figure 4** Rarefaction curves showing the number of observed species *per* marine sample sequenced with Illumina technology.

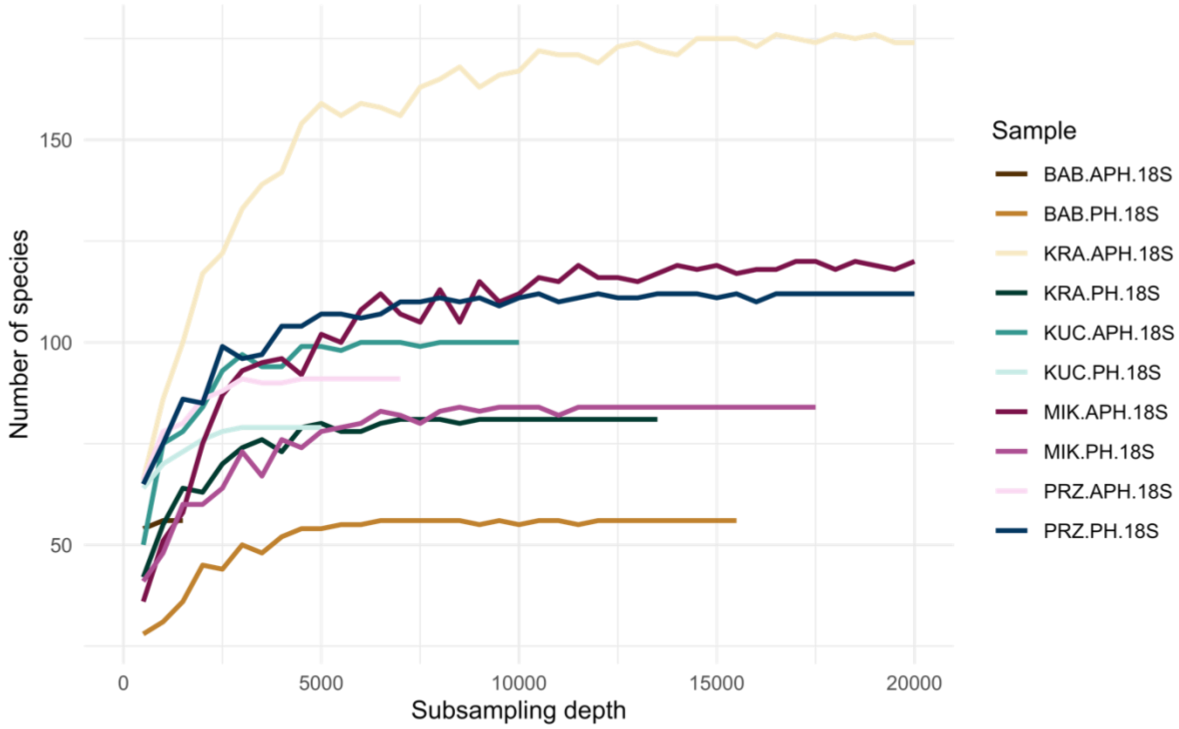


**Supplementary Figure 5** Rarefaction curves showing the number of observed species *per* freshwater sample sequenced with Nanopore technology.

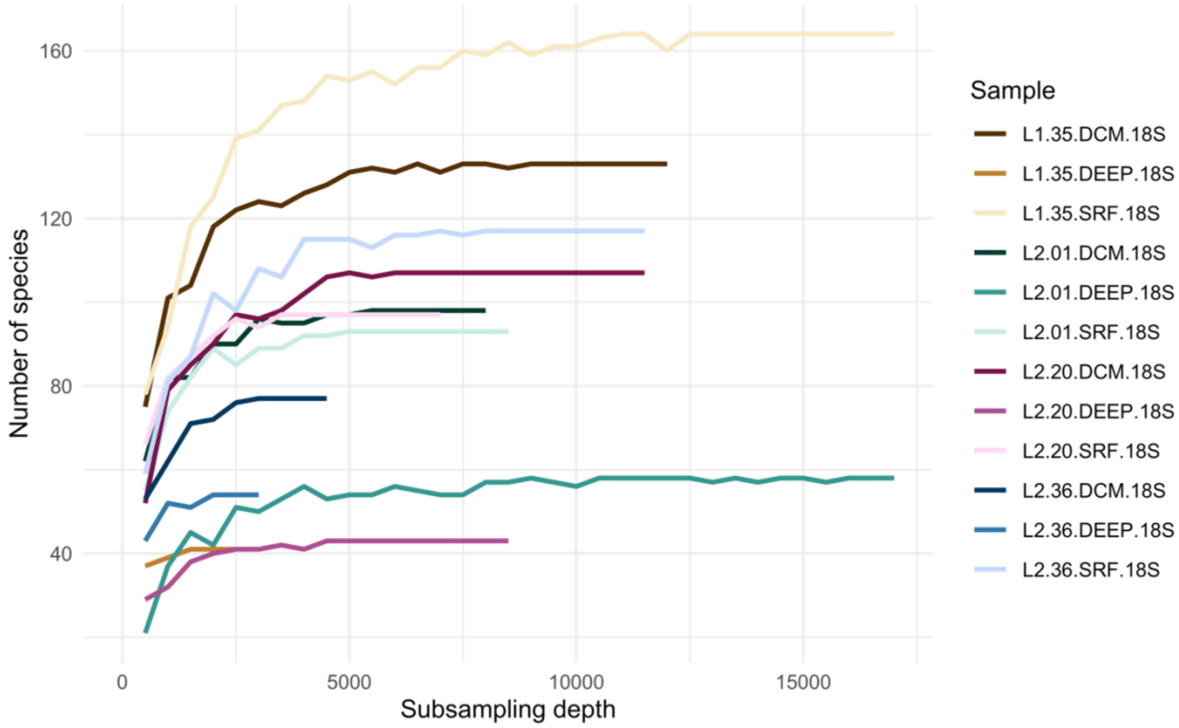


**Supplementary Figure 6** Rarefaction curves showing the number of observed species *per* marine sample sequenced with Nanopore technology.
